# Transcriptome Reprogramming Associated with Autoimmunity Reveals Key Points of the Growth-Defence Trade-off in *Arabidopsis thaliana*

**DOI:** 10.1101/2025.06.30.662188

**Authors:** Donghui Hu, Jinge Wang, Rachelle R.Q. Lee, Zezhao Su, WangSheng Zhu, Eunyoung Chae

**Affiliations:** Department of Biological Sciences, National University of Singapore, 117558, Singapore; China Key Laboratory of Pest Monitoring and Green Management, MOA, and College of Plant Protection, China Agricultural University, Beijing, 100193, China; Department of Biology, University of Oxford, Oxford, OX1 3RB, UK

**Author notes:** Corresponding author: Eunyoung Chae.

**Keywords:** plant immunity, growth-defence trade-off, autoimmunity, *ADR1* helper NLRs, immune regulation, *Arabidopsis thaliana*

## Abstract

Growth and defence are inversely correlated processes requiring balance for optimal fitness. While each is well studied, their regulatory trade-offs are less understood. Using *DANGEROUS MIX* (*DM*) autoimmune plants with stunted growth in *Arabidopsis thaliana*, we investigated the balancing act. Transcriptome analysis of three *DM* cases and pathogen-treated seedlings identified two major modules representing defence and growth. These core modules, comprising 4,712 genes (∼17% of the transcriptome), were sufficient for projecting the trade-off across immune transcriptome meta-datasets. Removing all three *ADR1s*, the helper NLRs, reversed the expression patterns of both modules and shifted the trade-off balance. *ADR1s* had a stronger impact on growth-associated RLK genes than on defence genes. Autoimmune plants also showed greater chromatin accessibility changes in growth loci than in defence loci. Our findings suggest an unexpected mechanism where immune signalling, partly mediated by *ADR1s*, actively suppresses growth genes, providing a strategy to regulate the growth-defence trade-off.

## Introduction

Plants navigate complex microbial landscapes through immune networks that exhibit a funnel-shaped architecture—diverse upstream inputs converging on shared signalling hubs to produce coordinated physiological outputs (Lapin et al., 2020; Pruitt et al., 2021; Lee and Chae, 2025). Central to successful defence is rapid cellular reprogramming that involves extensive gene expression changes. Both pattern-triggered immunity (PTI) and effector-triggered immunity (ETI) result in massive but overlapping transcriptome changes (Tsuda et al., 2009; Tsuda and Katagiri, 2010; Buscaill and Rivas, 2014; Tsuda and Somssich, 2015), despite the fact that immune triggers and the recognizing receptors display striking diversity. PTI ensues through the broad-spectrum recognition of pathogen/microbe-associated molecular patterns (PAMP/MAMP) via cell-surface receptors, including Receptor-Like Kinases (RLKs) and Receptor-Like Proteins (RLPs), whilst ETI engages intracellular nucleotide-binding leucine-rich repeat (NLR) receptors to detect specific pathogen effectors (Dangl and Jones, 2001; van der Hoorn and Kamoun, 2008; Boutrot and Zipfel, 2017; Saijo et al., 2018). This transcriptome reprogramming often associates with common plant immune responses, including calcium influx, reactive oxygen burst, cascaded MAP kinase activation, defense hormone synthesis such as salicylic acid (SA) and jasmonic acids (JA). Recent studies have validated PTI-ETI interplays, which has long been conceptualized based on blurred boundaries (Thomma et al., 2011; Cui et al., 2015; Ngou et al., 2021; Yuan et al., 2021). These highlight the central role of signaling hubs in orchestrating robust defence mechanisms (Schulze et al., 2022; Yang et al., 2022). Furthermore, plant immune responses share non-trivial transcriptional overlap with abiotic stress responses, suggesting cross-talk in stress responses (Jacob et al., 2018; Saijo and Loo, 2020; Yang et al., 2021)

While the overlap is evident, ETI, involving the activation of NLRs, often exerts acute and self-destructive responses, leading to tissue collapse through the hypersensitive response (HR), an immunological cell death in plants (REF). Two main classes of NLRs, CC-NBS-LRR (CNL) and TIR-NBS-LRR (TNL), form resistosomes upon effector recognition and activate immunity through distinct mechanisms (Huang et al., 2022; Jia et al., 2022; Förderer et al., 2022b). While CNL resistosomes can function as calcium channels to directly induce cell death (HR) (Bi et al., 2021; Förderer et al., 2022b), TNL signaling relies on small signaling molecules generated by the NADase activity of their TIR domains, of which holoenzyme activities are assembled in the TNL resistosomes (Huang et al., 2022; Jia et al., 2022). Upon the small molecule binding to the EDS1-containing signaling hub, another class of NLRs containing RPW8 domains (RNLs), so called helper NLRs, are tethered to the EDS1-complex to execute full effect (REF). Phylogenomic analyses have revealed that RNL diversity extends far beyond the well-characterised ADR1 and NRG1 families, encompassing multiple independent evolutionary lineages in gymnosperms with distinct distribution patterns across plant taxa (Andolfo et al., 2019; Qin et al., 2024). Unlike the highly diverse and rapidly evolving sensor NLR repertoires driven by pathogen selection pressures, helper NLRs exhibit greater evolutionary conservation and more ancestral phylogenetic patterns (Van de Weyer et al., 2019; Liu et al., 2021). This conservation reflects their fundamental role as central signalling hubs that must maintain compatibility with diverse upstream sensors while coordinating complex downstream responses.

The ADR1 family represents one well-studied subset within the broader RNL group. In *Arabidopsis thaliana* (herein Arabidopsis), three ADR1s (*ADR1*, *ADR1-L1*, and *ADR1-L2*) function as signal transducers downstream of sensor NLRs (Bonardi et al., 2011; Dong et al., 2016; Wu et al., 2021; Wang et al., 2021), forming complexes with EDS1-PAD4 to amplify immune signalling and orchestrate transcriptional reprogramming (Lapin et al., 2019). Recent studies demonstrate that ADR1s are more ubiquitous across angiosperms compared to NRG1s (Liu et al., 2021). NRG1s are typically associated with TNL-mediated immunity and are absent in lineages where TNLs are reduced (Liu et al., 2021). The broader distribution of ADR1s suggests that ADR1s serve as core, conserved components of plant immune networks with fundamental roles in immune signaling. However, this conserved function also creates potential vulnerabilities—immune network convergence points such as EDS1 can become single points of failure susceptible to dysregulation in hybrid contexts (Dongus and Parker, 2021; Lee and Chae, 2025).

This convergent organisation becomes important in cases of autoimmunity, where dysregulated immune signalling leads to constitutive defence activation at the expense of normal growth and development. The *DANGEROUS MIX* (*DM*) system, established from the studies of hybrid necrosis in Arabidopsis, demonstrates how incompatible NLR combinations can exploit these immune network vulnerabilities, where combinations of incompatible NLR pairs from different accessions trigger not only constitutive immune activation but also severe growth penalties (Bomblies and Weigel, 2007; Chae et al., 2014). Hence, the *DM* system offers a powerful model for studying how the shift in growth-defence balance is achieved through different NLR signaling. Well-characterised examples include *DM1*^Uk-3^-*DM2d*^Uk-1^, where heteromeric TNL oligomer formation activates EDS1-dependent signaling (Bomblies et al., 2007; Tran et al., 2017); *DM10*^TueScha-9^-*DM11*^Cdm-0^ involving a truncated TNL (*DM10*) with extremely severe necrosis (Barragan et al., 2021);. *DM6*^Lerik1-3^-*DM7*^Fei-0^ of which signaling is dependent on the P-loop of the CNL, DM6/RPP7b (Barragan et al., 2019; Li et al., 2020). Of particular interest is that the *DM9* cases are due to incompatibilities between the alleles of the same locus, *ACCELERATED CELL DEATH 6* (*ACD6*) (Todesco et al., 2014; Świadek et al., 2017; Zhu et al., 2018). Natural allelic variation of *ACD6* as well as its mutant studies have marked its central position in regulating the growth-defence trade-off (Lu et al., 2003a; Todesco et al., 2010). These growth penalties were also extensively described in Arabidopsis autoimmune mutants. such as *acd6-1*, *bak1-4 serk4-1*, and *ssi2* (Alcázar et al., 2009; de Oliveira et al., 2016; Yang et al., 2016; Atanasov et al., 2018; Zhang et al., 2019; Yang et al., 2020).

Whether immune signalling actively suppresses growth through dedicated circuits, rather than simple resource competition, remains unclear (Huot et al., 2014; He et al., 2022). A fundamental challenge is to determine whether this trade-off can be decoupled from complex downstream of immune activation.

Here, we combine multiple transcriptomic and chromatin accessibility analyses of *DM*-triggered autoimmunity in WT and *adr1 triple* knockout (KO). By eliminating ADR1 function, we reveal their dual role in enhancing immune responses and actively suppressing growth-related genes. Integrating RNA-Seq with ATAC-Seq data, we identify the transcription factors and chromatin remodelling events mediating this bifurcated response, establishing ADR1s among master regulators of the growth-defence trade-off and revealing potential targets for engineering.

## Results

### Extensive Transcriptomic Reprogramming and Enhanced NLR Transcription in *DMs*

To investigate autoimmune signalling triggered by different NLR types, we selected the DM1^Uk-3^-DM2d^Uk-1^ case signaling through the canonical TNL pathway through EDS1 (Tran et al., 2017) and the DM6^Lerik1-3^-DM7^Fei-0^ case involving the resistosome formation of CNL DM6/RPP7with a cognate DM7/RPW8 partner (Barragan et al., 2019; Li et al., 2020). We assembled these *DM* pairs into single constructs to establish stable Col-0 transformants, which exhibited characteristic autoimmune phenotypes (**Figure 2A**). The homozygous transformants displayed severe growth arrest, developing only rudimentary true leaves beyond cotyledons, with *DM1-DM2d* presenting more acute symptoms characterized by chlorosis. RNA-Seq was performed on aerial tissues of homozygous transformants expressing *DM1-DM2d* or *DM6-DM7*, alongside Col-0 at 12 and 14 DAS, respectively. Integration with the published *DM10-DM11* data, involving a truncated TNL encoded by *DM10* (Barragan et al., 2021), revealed parallel transcriptomic shifts across these configurations.

Principal component analysis (PCA) highlighted divergence between *DMs* and WT backgrounds, with PC1 explaining 62.3% variance. Transcriptomic deviations were more pronounced in *DM10-DM11* and *DM1-DM2d* than in *DM6-DM7* (**Figure 1A**). Collectively, these *DM* instances exhibited extensive gene expression changes, with 4,697 up-regulated and 5,625 down-regulated genes across all cases under stringent cutoffs (log2FC > 2 or < −2, adjusted p-value < 0.05), representing approximately 38% of detected genes. Among these, 1,039 genes were commonly up-regulated and 1,126 commonly down-regulated across all three *DM* cases (**Figure 1B**). TNL-associated *DMs* (*DM10-DM11* and *DM1-DM2d*) shared more DEGs with each other than with CNL/RPW8-associated *DM6-DM7*. GO analysis of up-DEGs revealed enrichment in “programmed cell death”, “response to salicylic acid”, and “defence response”, indicating enhanced immune states. Down-DEGs were associated with “developmental process”, “lipid metabolism”, and “response to auxin”, suggesting reduced growth (**Figure S1A-S1B**). Photosynthesis-related GO terms were notably prevalent in down-DEGs of *DM10-DM11* and *DM1-DM2d*.

**Figure 1.**
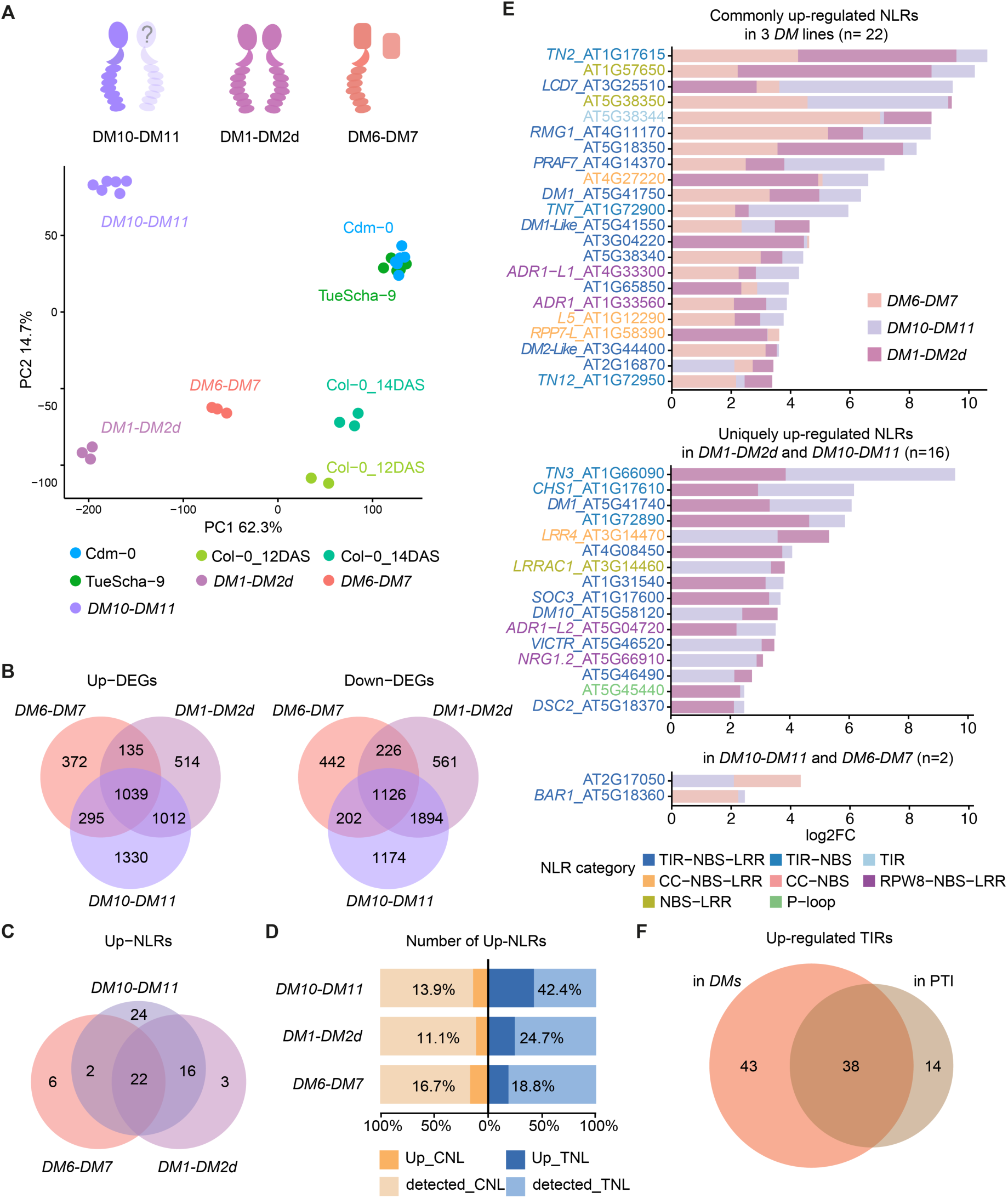
mRNA-Seq analysis unveils extensive transcriptomic reprogramming and augmented NLR transcription in three autoimmune lines. (**A**) Upper panel: Scheme of DMs. DM10 is identified as TNL, DM1-DM2d is TNL pair. DM6-DM7 is a pair of CNL and RPW8. Bottom panel: Principal component analysis (PCA) plot of three DMs RNA-Seq samples. PC1 separated the DMs from WT, which is DM10-DM11 vs two parents, TueScha-9 and Cdm-0; DM1-DM2d vs Col-0 12 DAS; DM6-DM7 vs Col-0 14 DAS. (**B**) Venn Diagram of up-regulated genes in three autoimmune lines, DM10-DM11, DM1-DM2d and DM6-DM7, versus their WT control, with log2FoldChange (log2FC) > 2, adjusted p-value < 0.05. Up-DEGs, up-regulated differentially expressed genes (DEGs). Down-DEGs, up-regulated differentially expressed genes (DEGs). (**C**) Venn Diagram of up-regulated NLRs in three DMs with log2FC > 2. (**D**) the ratio of up-regulated CNL and TNL in the total detected CNL/TNL. (**E**) Bar plot of up-regulated NLRs. Three DMs have overlapped bars. Labels of NLRs are colour-coded. log2FC are calculated as each DMs vs their own control. (**F**) Venn diagram of up-regulated TIR-containing genes (log2FC > 1 in any DMs vs WT or in elicitor triggered PTI vs mock control). Elicitor-triggered PTI data from Tian et al., 2021 that applied log2FC > 1. A relaxed fold change was applied to our dataset for comparison.

A notable feature was widespread NLR up-regulation (**Figure S1C**), with 73 of 163 NLR genes significantly up-regulated in at least one *DM* case and 22 NLRs consistently elevated across all three (**Figure 1C**). TNL-involving *DMs* induced more NLR genes, with *DM1-DM2d* and *DM10-DM11* sharing 16 up-regulated NLRs whilst *DM10-DM11* alone included an additional 24. This response was enriched with TNL genes in TNL-involving *DM* pairs, with *DM10-DM11* up-regulating 36 of 85 detected TNLs (42.4%) (**Figure 1D**). TNL gene induction correlated with phenotypic severity: *DM10-DM11* showed highest induction (42.4%), followed by *DM1-DM2d* (24.7%) and *DM6-DM7* (18.8%). CNL up-regulation was comparatively modest and consistent across all cases (13.9%, 11.1%, and 16.7% in *DM10-DM11*, *DM1-DM2d*, and *DM6-DM7*, respectively) (**Figure 1D**).

Among the 22 commonly up-regulated NLRs, TIR-containing genes were enriched, including both full-length TNLs and truncated TNs lacking LRR domains (**Figure 1E**). From 136 TIR-containing genes in *A. thaliana* (Wan et al., 2019), 81 were up-regulated in at least one *DM* condition, whilst Tian et al., 2021 reported 52 up-regulated by PTI-triggering elicitors. *DM*-triggered autoimmunity and PTI responses shared 38 up-regulated TIR genes (**Figure 1F**, **Table S1**), suggesting TIR gene upregulation is common between these distinct immune triggers. Helper RNLs constituted a commonly upregulated group; two *ADR1s* family members, *ADR1* and *ADR1-L1*, appeared within commonly activated NLRs, whilst *ADR1-L2* and *NRG1.2* showed heightened activation in TNL-associated cases.T his concerted NLR activation suggests a hierarchical switching mechanism activating numerous NLRs simultaneously upon initial immune triggering. Given that *ADR1* overexpression via activation tagging confers autoimmune phenotypes (Grant et al., 2003) and helper NLRs relay immune signalling, *ADR1* and its homologues are prime candidates for mediating NLR upregulation downstream of the *DM*-NLR immune switch.

### *DM6-DM7* and *DM1-DM2d* Autoimmunity are Mitigated by *adr1 triple* Knock-Out

To test dependency of *DM*-mediated immune signalling on *ADR1s*, we designed a multiplexed CRISPR-Cas9 strategy that simultaneously targets *ADR1*, *ADR1-L1* and *ADR1-L2* and generated their triple knock-out (*adr1 triple*) in two *DM* lines and Col-0 (**Figure S2**). The *adr1 triple* mutation in *DM1-DM2d* markedly improved growth and development, evidenced by restoration of green leaves from chlorosis, though autoimmune symptoms persisted (**Figure 2A and S3**). Hemizygous *DM1-DM2d* plants with *adr1 triple* showed increased rosette size (**Figure 2A**). While the mutation did not alter the appearance of homozygous *DM6-DM7* plants, it modestly increased the size of hemizygous ones (**Figure 2B and S3**). The enhanced growth observed in both Col-0 and WT-segregants from hemizygous *DM*s suggests that *ADR1s* modulate the growth-defence trade-off (**Figure 2C**). This growth improvement persisted into later developmental stages. *DM6-DM7 adr1 triple* and Col-0 *adr1 triple* maintained larger size than their respective backgrounds until flowering, whereas *DM1-DM2d adr1 triple* plants, though initially larger, developed distinct autoimmune phenotypes, such as twisted leaves, from 18 DAS onwards (**Figure S3**). Quantitative PCR (qPCR) analysis confirmed a dose-dependent effect of *DM1-DM2d* construct copy number on the severity of autoimmune phenotypes. Homozygous plants exhibited more severe symptoms than hemizygous ones (**Figure S4**).

**Figure 2.**
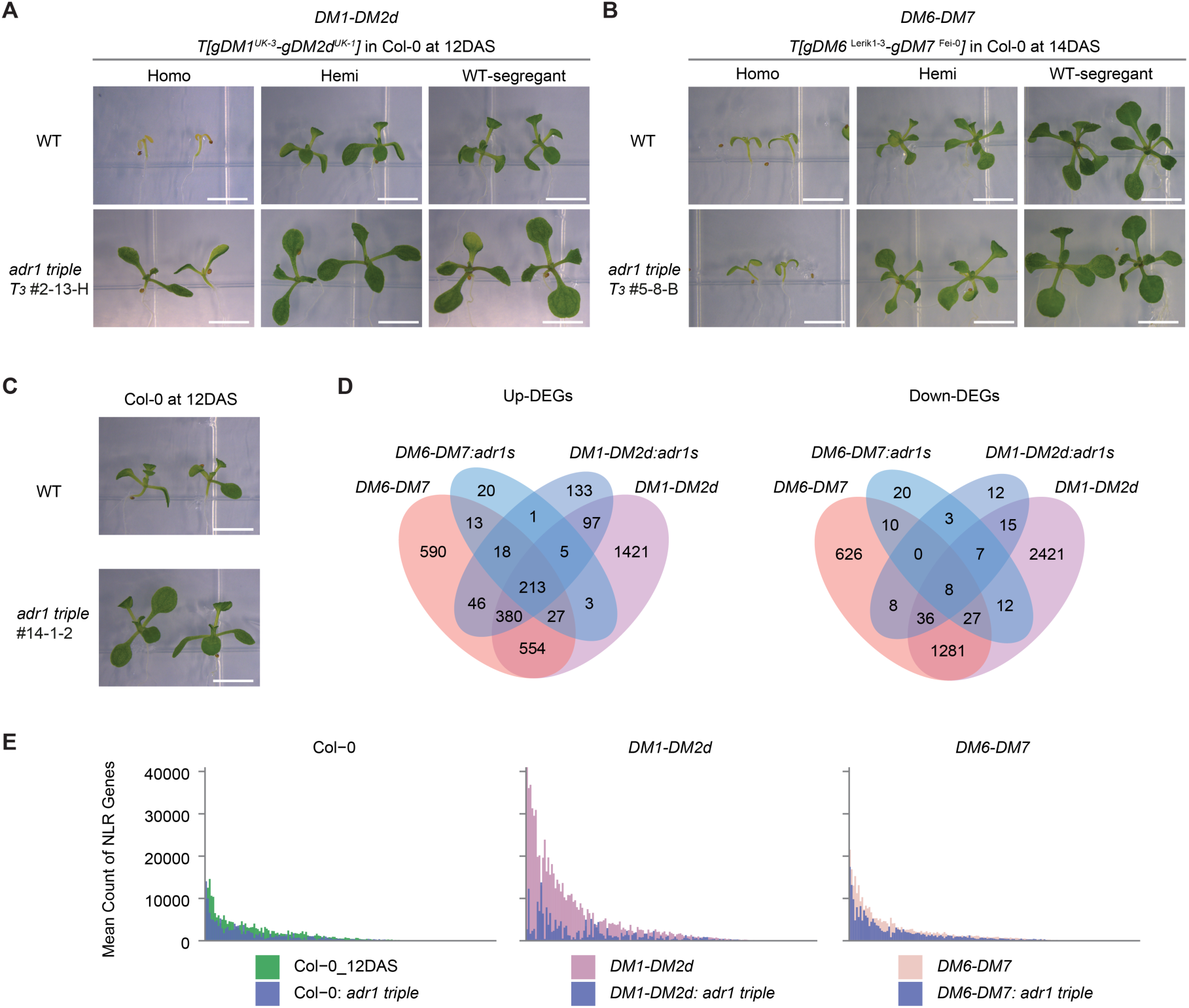
Phenotypes of *DM6-DM7* and *DM1-DM2d* autoimmunity are partially rescued by *adr1 triple*. **(A)** *adr1 triple* in *DM1-DM2d* background. The autoimmunity phenotype observed in the *DM1-DM2d* gene pair is partially rescued by *adr1s triple*. “Homo” refers to homozygous plants, and “Hemi” refers to hemizygous plants. “WT-segregant” refers to segregated plants not carrying the transgenic insertion. The zygosity of the plants was confirmed by quantifying the copy number variation (CNV) of the transformed *DM1-DM2d* gene via qPCR. The photo was taken at 12 days after sowing (DAS). The scale bar represents 5 mm. **(B)** *adr1s triple* in *DM6-DM7* background. The autoimmunity phenotype observed in the *DM6-DM7* gene pair is also partially rescued *adr1s triple*. The photo was taken at 14 DAS. The scale bar represents 5 mm. **(C)** *adr1s* in the Col-0 background. This panel shows the effect of *adr1* knockout in the wild-type Col-0 background. The scale bar represents 5 mm. **(D)** Venn diagram of up-regulated and down-regulated genes in *DM6-DM7*, *DM1-DM2d* and their respective *adr1s triple* mutants. This diagram provides an overview of the gene expression changes in response to *adr1 triple* in different backgrounds. The significance is defined by | log2FC | > 2 and adjusted p-value padj < 0.05. **(E)** NLRs expression profile in different backgrounds. The plot is faceted by the background, showing the expression profile of NLRs in Col-0, *DM1-DM2d*, and *DM6-DM7* backgrounds.

We extended RNA-Seq to include the *adr1 triple* mutants in these backgrounds to reveal transcriptional mechanisms underlying phenotypic recovery. RC1 (28% variation) of rotated PCA separated autoimmune *DM* lines from WT, with *adr1 triple DM* mutants positioned intermediately, indicating partial transcriptome restoration. Col-0 *adr1s* samples clustered left of wild-type, suggesting reduced basal immunity (**Figure S5B**). The *adr1 triple* mutation dramatically reduced up-DEGs in *DM1-DM2d* (2700 to 893) and *DM6-DM7* (1841 to 300), with even more substantial decreases in down-DEGs in *DM1-DM2d* (3807 to 89) and *DM6-DM7* (1996 to 88) (**Figure 2D**).

Further analysis revealed that *adr1 triple* mutations reduced *DM*-induced NLR expression more significantly in *DM1-DM2d* (median read counts: 1513 to 459) than in *DM6-DM7* (761 to 512) (**Figure 2E**), correlating with the observed degree of phenotypic rescue (**Figure 2A and 2B**). Notably, *adr1 triple* in Col-0 also decreased NLR expression, paralleling the growth improvements seen in *DM6-DM7* (**Figure 2E**). This differential impact suggests context-dependent transcriptional modulation by ADR1s, with stronger influence on TNL-mediated autoimmunity pathways.

### Transcriptome Modules Shared Between Autoimmunity and Pathogen-induced Immunity

To investigate commonalities and differences between pathogen-treated effects and *DM*-induced autoimmunity, we employed a transient flood assay (Ishiga et al., 2011; Ishiga et al., 2017) with *Pseudomonas syringae* pv. *tomato* DC3000 (*Pto* DC3000) carrying the effector *AvrRpt2* or *AvrRps4* on Col-0 seedlings at equivalent developmental stages to the *DM* lines. Tissue necrosis emerged at 24 hpi and intensified markedly by 48 hpi, indicating progressive disease severity (**Figure S5A**). Time-series RNA-Seq experiments at 6h, 12h, and 20h captured temporal transcriptome dynamics alongside mock-treated controls. We identified a set of TIR-containing genes upregulated across all three conditions: PTI (Tian et al., 2021), *DMs*, and our flood assay, supposedly triggering both ETI and PTI (**Table S2**).

For meta-analysis, we combined these datasets with those from *DMs* in WT, the *adr1 triple* mutant backgrounds, and the *adr1 triple* line in Col-0 (**Figure S5B, Table S3**). Rotated PCA showed RC1 effectively distinguished *Avr*-treated samples from mock treatments, except for the earliest mock samples (6h) where flooding effects were apparent. Later mock treatments clustered at lower RC1 values, whilst pathogen-treated samples demonstrated time-dependent progression toward intermediate immune activation between WT and *DM* samples, with responses peaking at 20h. *AvrRpt2*-treated samples consistently exhibiting higher RC1 values than *AvrRps4* samples across all timepoints. The earlier necrosis onset with *AvrRpt2* (Saile et al., 2020) reflects differences in effector recognition of *AvrRps4* by TNL RPS4 versus *AvrRpt2* by CNL RPS2 (Bent et al., 1994; Hinsch and Staskawicz, 1996).

Weighted Gene Co-expression Network Analysis (WGCNA) further analysed the metadata in an unsupervised manner (Langfelder and Horvath, 2008). WGCNA categorised the top 20% of most variable genes across all 71 RNA-Seq samples into 10 distinct modules (**Figure S5C-S5D, Figure 3B**). Two key modules were central to this transcriptome reprogramming: the common immune response module MEruby (n=2490) and the suppressed growth module MEblue (n=2222) alongside a smaller but related MEpaleblue module (n=234) (**Figure 3B**).

**Figure 3.**
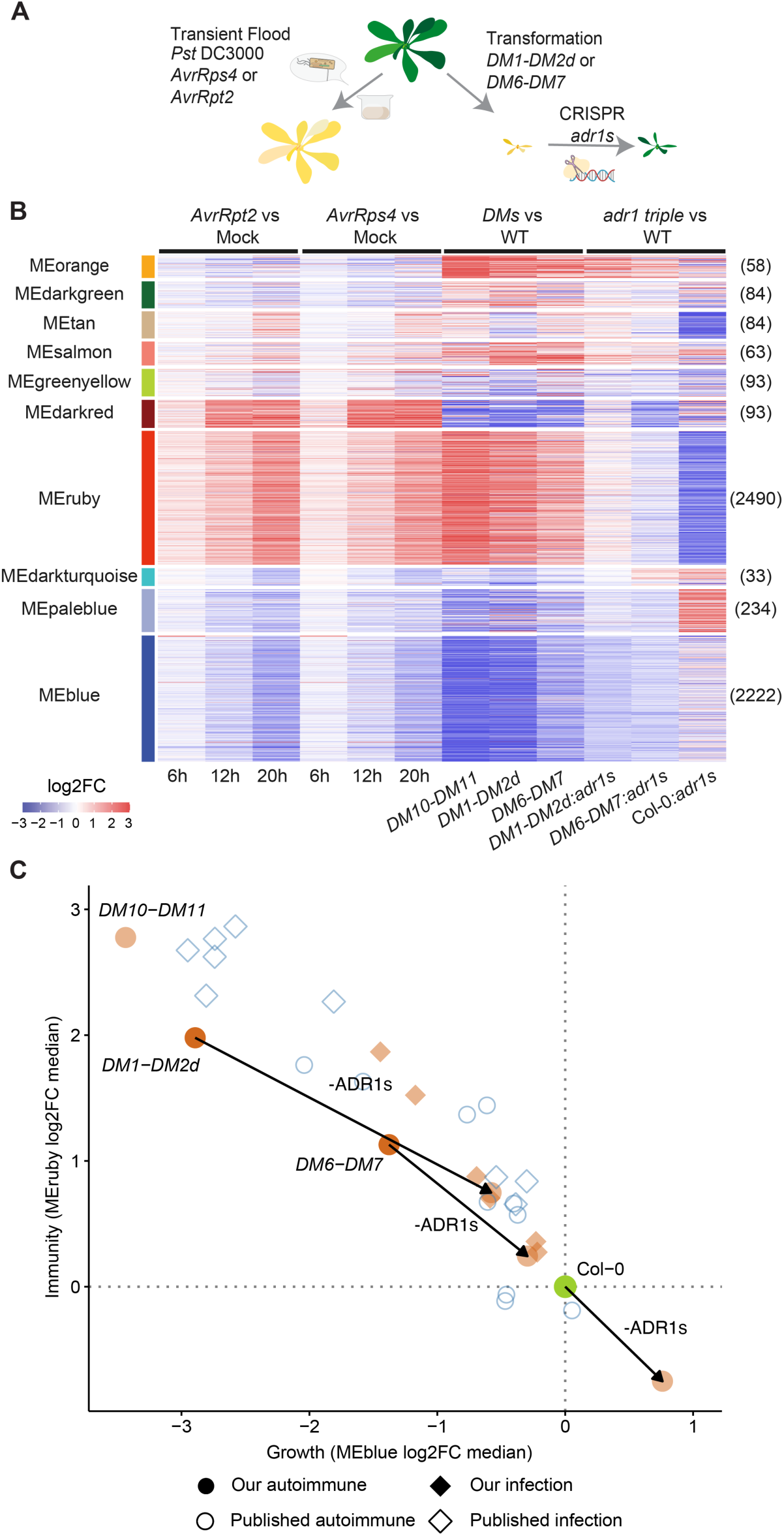
WGCNA analysis reveals shared and distinct transcriptional modules between pathogen-induced immunity and autoimmunity. **(A)** Schematic representation of experimental design. Wild-type Col-0 seedlings were subjected to transient flood treatment with *Pst* DC3000 expressing either *AvrRps4* or *AvrRpt2* effectors. In parallel, Col-0 plants were transformed with either *DM1-DM2d* or *DM6-DM7* pairs, followed by CRISPR-mediated knockout of *ADR1* genes (*adr1* triple mutant). **(B)** Heatmap showing log2FC values of gene expression across different experimental conditions and genotypes. WGCNA analysis identified distinct co-expression modules (shown on y-axis with different colours). Numbers in parentheses indicate the number of genes in each module. Module heights in the heatmap are square root-transformed to improve visibility of smaller modules. The analysis compares four conditions: *AvrRpt2* vs Mock, *AvrRps4* vs Mock (at 6h, 12h, and 20h), *DMs* vs WT, and *adr1* triple mutant vs WT. Red indicates upregulation, and blue indicates downregulation relative to controls. **(C)** Quantitative visualisation of the growth-defence trade-off across diverse experimental conditions. Left panel: Scatter plot showing MEruby (immunity, y-axis) versus MEblue (growth, x-axis) log₂FC medians across all samples analysed. Each point represents a specific experimental condition or genotype. The linear relationship between growth suppression and immunity activation demonstrates that *ADR1s* function as central modulators coordinating this antagonistic relationship. Col-0 serves as the reference point positioned at the origin, with pathogen treatments and autoimmune conditions progressing along a consistent negative correlation trajectory, suggesting a universal regulatory mechanism balancing these competing physiological demands. Right panel: Focused view highlighting the transition from autoimmune *DM* states to their *adr1* triple rescue mutants. Lines connect *DM* lines or Col-0 to their corresponding *adr1* triple mutants, revealing consistent shifts towards enhanced growth and reduced immunity when *ADR1s* are knocked out.

The MEruby module showed progressive upregulation in both *Pst*-treated Col-0 and *DM* lines, with *AvrRpt2* eliciting a slightly quicker and stronger response than *AvrRps4*. GO enrichment analysis revealed upregulation of biotic stress response genes, including immune system processes, defence response, and programmed cell death (**Figure S6**). Conversely, MEblue exhibited progressive downregulation, with GO analysis showing enrichment for vital growth and maintenance functions like photosynthesis and plastid organisation. The severity of transcriptomic bifurcation correlated with autoimmunity intensity, with TNL-associated *DM10-DM11* and *DM1-DM2d* showing more extreme patterns than the CNL/RPW8 pair *DM6-DM7* (**Figure 3B**). Notably, the *adr1 triple* mutation moderated these transcriptomic extremes in all *DM* backgrounds, reducing upregulation in MEruby and lessening downregulation in MEblue. Intriguingly, the Col-0 *adr1 triple* exhibited an inverse pattern— downregulation in MEruby and upregulation in MEblue compared to Col-0 WT— revealing that ADR1s moderate growth-immunity balance even without obvious triggers. The MEpaleblue module showed similar expression patterns to MEblue but with stronger upregulation in the Col-0 *adr1s triple*, indicating these genes (enriched for metabolic and developmental functions) are effectively suppressed by ADR1s in WT (**Figure S6D**).

Two other modules with distinct patterns were identified: the prolonged immune response MEorange and the pathogen-induced response MEdarkred (**Figure 3B**). (**Figure 3B**). The MEorange, uniquely upregulated in *DMs* but not in pathogen-treatment samples, reflects physiological responses to prolonged autoimmunity. It contains immune components, including LRR-RLPs (*RLP34*, *RLP37*, *RLP47*), the calcium channel *CYCLIC NUCLEOTIDE-GATED CHANNEL 11* (*CNGC11*), *ACD6* and receptor-like kinases *CRK4* and *CRK5*, with GO terms related to general defence responses, programmed cell death and abscisic acid response (**Figure S6B**). In contrast, the MEdarkred module is uniquely upregulated during pathogen treatment and downregulated in *DMs*. It is enriched with jasmonic acid (JA)-responsive genes, including *PLANT DEFENSIN* genes (*PDF1.2*, *PDF1.2c*, *PDF1.3*) and *VEGETATIVE STORAGE PROTEIN* genes (*VSP1*, *VSP2*). This suggests active JA signalling during the flood assay compared to *DM* conditions, consistent with coronatine effects from Pst (Zheng et al., 2012).

For cross-validation, we examined WGCNA modules profiles across diverse published transcriptome datasets using MEruby and MEblue, which are ∼17% of transcriptome (**Figure S7, Table S4**), including time-course studies of bacterial effector responses (*AvrRpm1*, *AvrRpt2* or *AvrRps4* vs *EV* by infiltration) (Saile et al., 2020), an estradiol-inducible *AvrRps4* system (Ngou et al., 2021), and several autoimmune mutants: *acd6-1* (Zhang et al., 2019), *bak1-4 serk4-1* (de Oliveira et al., 2016), *ka120* (Jia et al., 2021), *ssi2-1, ssi2-2 and ssi2-3* (Yang et al., 2016), and *hos15-4* (Yang et al., 2020). Another genetic incompatibility study comparing Ler/Kas-2 NIL with Kas-2 and its suppressor *sulki1-8* (Alcázar et al., 2009; Atanasov et al., 2018) is also included. Meta-analysis across these diverse datasets faithfully reproduced the MEruby and MEblue module expression patterns identified in our studies. This remarkable conservation across independent experimental systems confirms that these modules represent universal transcriptional signatures of plant immune responses. Meanwhile, the gene family analysis identified that the majority of NLRs, TIR-containing genes are commonly up-regulated in autoimmunity, while Col-0 *adr1s* showed a mainly down-regulation of them (Figure S8A-S8B).

Furthermore, we documented a striking negative linear correlation that provides compelling molecular evidence for the growth-defence antagonism using MEruby and MEblue modules (**Figure 3C**). The gradient of this negative correlation appears consistent across datasets: As the reference sample, Col-0 is anchored at the origin, with increasingly activated immunity cases. progress further along the negative correlation trajectory. The *DM1-DM2d* line exhibits the most extreme trade-off, reflecting its severe autoimmunity. This correlation holds across diverse datasets: pathogen exposure time courses (our flood assay and Saile’s experiments, **Figure S7B**) and various autoimmune contexts (our *DM* lines alongside previously reported mutants including *acd6-1*, *bak1-4 serk4-1*, *ssi2* alleles, *hos15-4*, and Ler/Kas-2 NIL with *sulki1-8,* **Figure S7C**) (Alcázar et al., 2009; de Oliveira et al., 2016; Yang et al., 2016; Atanasov et al., 2018; Zhang et al., 2019; Yang et al., 2020). These diverse systems consistently shift away from baseline growth toward enhanced defence along the same cline, suggesting a universal regulatory mechanism balancing these competing physiological demands. In opposite, *adr1 triple* mutants shift autoimmune transcriptomes toward baseline along this same regulatory axis, indicating ADR1s function as central modulators of this antagonistic relationship. Removing *ADR1s* enhances growth in *DMs* backgrounds whilst reducing baseline immunity in WT.

WGCNA module patterns showed remarkable conservation across independent datasets, confirming MEruby and MEblue as universal markers of immune activation and growth suppression. Cross-validation across diverse experimental systems revealed MEorange as a specific indicator of prolonged autoimmunity, whilst the flood assay uncovered temporal dynamics in the MEdarkred module not apparent in traditional infiltration studies (**Figure 3B, Figure S7A**). The striking linear correlation between growth suppression and defence activation across all datasets (**Figure 3C, Figure S7B-C**) provides compelling molecular evidence for the growth-defence trade-off, with *adr1 triple* mutants consistently shifting autoimmune transcriptomes towards baseline, establishing ADR1s as central modulators of this antagonistic relationship.

### The Role of *ADR1s* in Maintaining Basal Expression of NLRs, RLPs and defence RLKs and in Suppressing Expression of Developmental RLKs

Our meta-analysis revealed *ADR1*s’ role in regulating both defence and growth-related gene modules (**Figure 3B**). Transcriptome profile from the *adr1 triple* mutant showed significant reversal across multiple modules (MEruby, MEtan, MEpaleblue and MEblue), prompting us to investigate *ADR1*-dependent signalling.

Of the 58 NLR genes in our WGCNA, most (n=37) belonged to the MEruby (common immune responses), while only three appeared in growth-suppressed modules (MEblue and MEpaleblue) (**Figure 4**). Nine NLRs in the MEsalmon module showed slight upregulation in both *DMs* and pathogen treatments. This distribution confirms previous findings of NLR expression escalation during biotic stress (Yang et al., 2016). The skewed distribution pattern, revealed through WGCNA filtering of our RNA-Seq across autoimmunity and pathogen treatment, suggests a general regulatory mechanism that amplifies defence through collective NLR upregulation. Notably, 27 TIR-containing genes were among the highly upregulated NLRs (log2FC > 2) (**Figure 4**), consistent with our *DM* analysis (**Figure 1E**). While MEruby NLRs are strongly induced in *DMs*, introducing *adr1 triple* mutations only moderately reduced their upregulation, suggesting the existence of an additional regulatory circuit. Interestingly, basal expression of most NLRs (except *ZAR1*, AT1G31540 and *DM10*) was reduced in the *adr1 triple* Col-0 mutant compared to Col-0, indicating *ADR1s* maintain baseline NLR expression. NLRs in smaller modules such as MEsalmon showed heterogeneous *ADR1* dependency (**Figure 4**).

**Figure 4.**
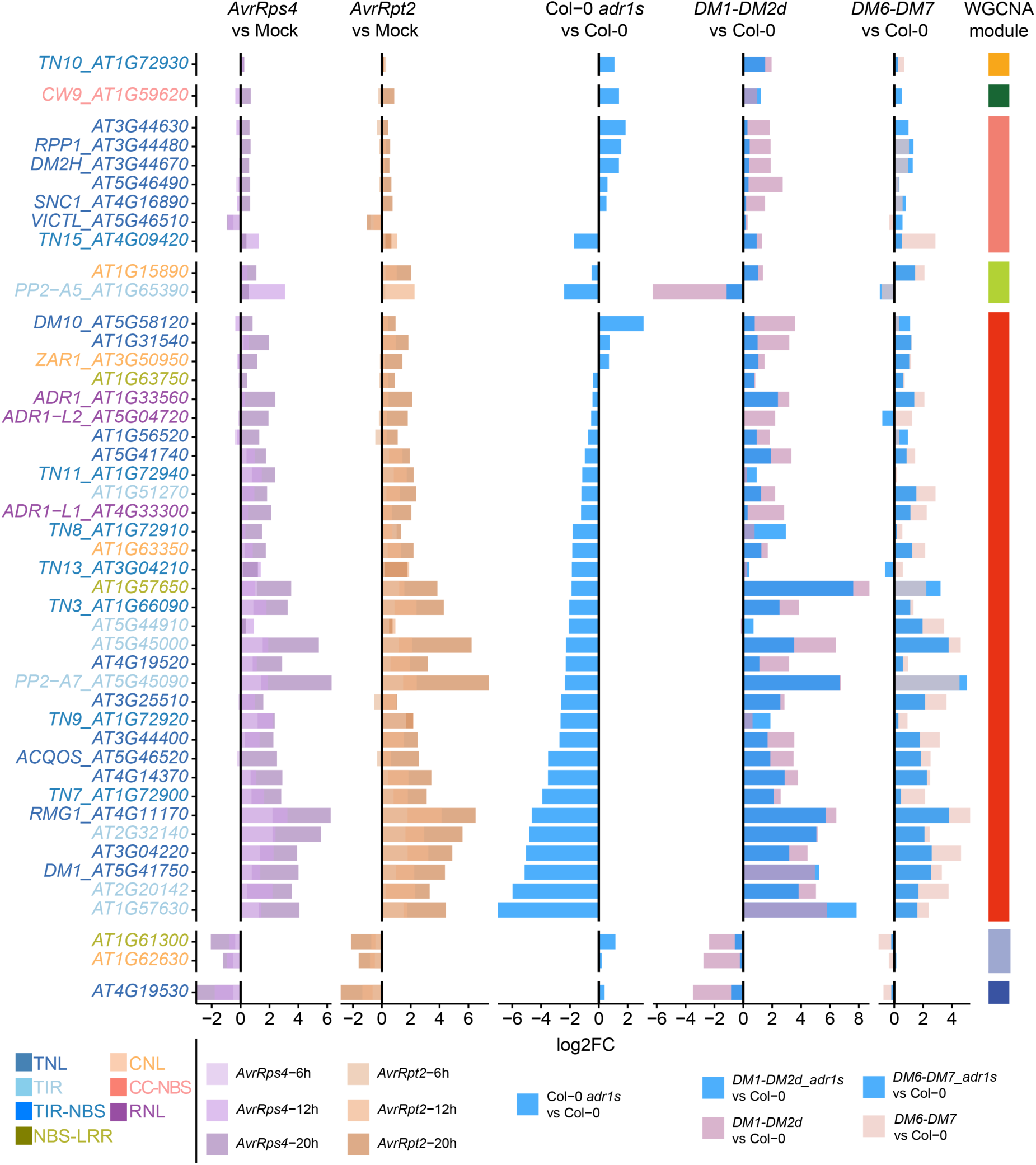
Differential expression of NLR genes post-pathogen treatment and in autoimmune backgrounds. *AvrRps4* and *AvrRpt2* are compared with Mock treatment at the same hpi. Col-0 *adr1s* is compared with Col-0 at 12 DAS. *DM1-DM2d* and *DM1-DM2d adr1s* are both compared with Col-0 at 12 DAS. *DM6-DM7* and *DM6-DM7 adr1s* are both compared with Col-0 at 14 DAS. The label of genes was colour coded.

Similar to NLRs, LRR-RLPs and some LRR-RLKs show upregulation and are primarily categorised in the MEruby module (10 RLPs out of 15 and 32 RLKs out of 88) (**Figure 5, Figure S8C-S8D**). These RLPs and RLKs demonstrate a similar dependency on *ADR1*s for their expression as observed with NLRs. Several key examples include *RECEPTOR-LIKE PROTEIN 23* (*RLP23*), which binds to nlp20, a conserved fragment of bacterial virulence proteins called Necrosis and ethylene-inducing peptide 1-like proteins (Albert et al., 2015), and *FLG22-INDUCED RECEPTOR-LIKE KINASE 1* (*FRK1*), a marker for early flagellin responses (Asai et al., 2002). Like NLRs in the MEruby module, these defence-related RLKs show reduced expression in the *adr1 triple* mutant compared to Col-0.

**Figure 5.**
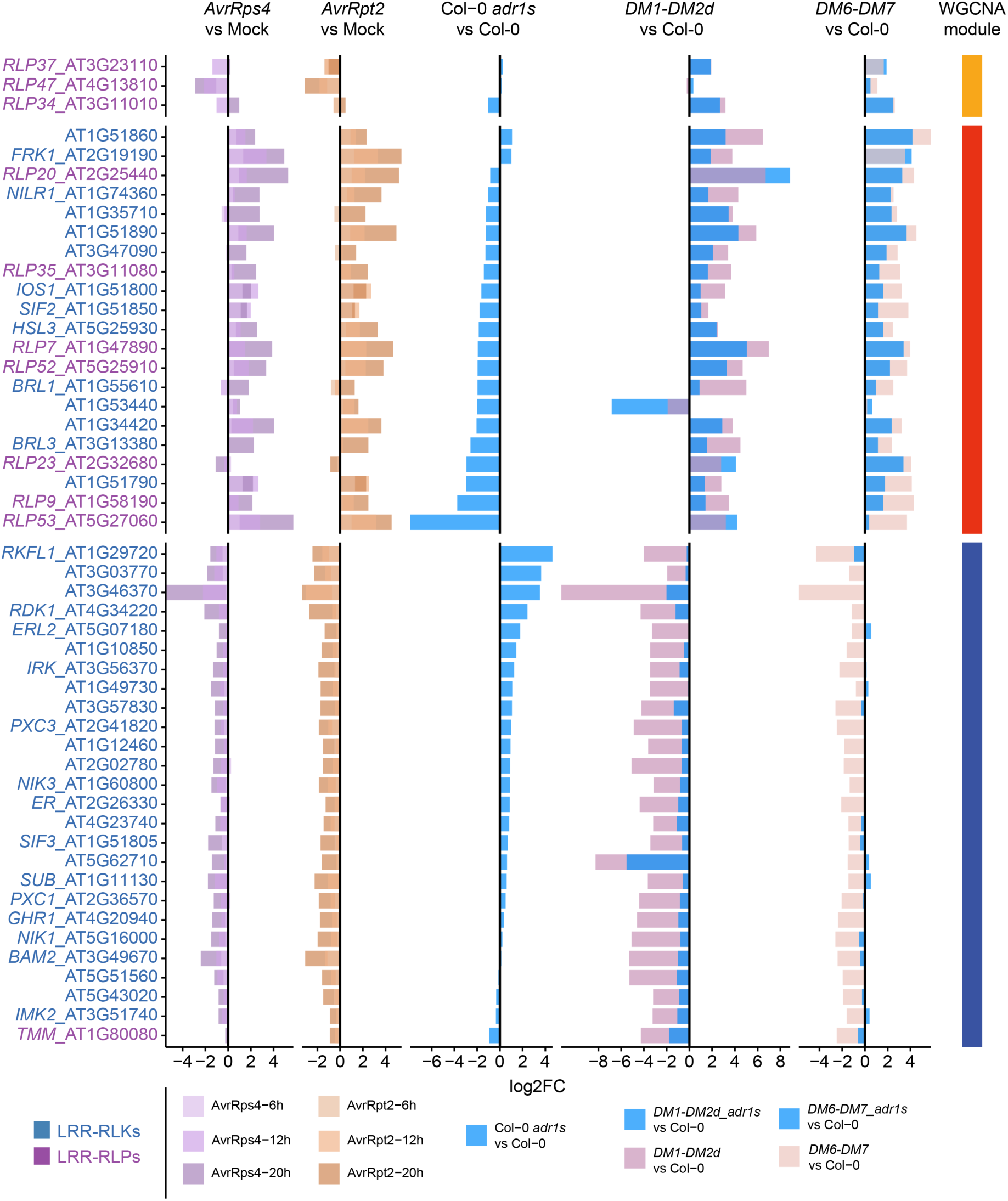
Differential Expression of LRR-RLP and LRR-RLK Genes Post-Pathogen Treatment and in Autoimmune Backgrounds. LRR-RLPs carry a deep orchid purple tone. LRR-RLKs are represented by a bold steel blue colour. To reduce the number of genes to plot, only genes that has log2FC > 3 in at least one comparison are included.

Interestingly, most RLKs (54 out of 88) were assigned to MEblue and exhibit an opposite pattern to NLRs and RLPs (**Figure 5, Figure S8C-S8D**). These MEblue RLKs are upregulated in the absence of *ADR1*s in Col-0, but dramatically downregulated in *DM1-DM2d*, reflecting severe growth penalties. This suppression is largely abolished by introducing the *adr1 triple* mutation in the *DM1-DM2d* background. The effect of *ADR1 triple* mutations on reversing growth gene suppression was significantly greater than their influence on reducing immune gene expression. This reveals *ADR1*s’ role in actively suppressing growth-related RLKs during immune responses, examples including *RECEPTOR DEAD KINASE* (*RDK1*), involved in seedling maturation (Kumar et al., 2017), *INFLORESCENCE AND ROOT APICES RECEPTOR KINASE* (*IRK*), which regulates root formation and cell division (Rodriguez-Furlan et al., 2022), and *ERECTA-LIKE 2* (*ERL2*), which promotes organ growth and flower development (Shpak et al., 2004) and so on.

The bifurcation in expression patterns between immune-related and growth-related genes in response to the mutation of ADR1s reveals their dual regulatory role. ADR1s contribute to maintaining basal expression of NLRs, defence-related RLPs and RLKs, while effectively suppressing growth-related RLKs during immune challenges. This dual function enhances pathogen detection and response capabilities while actively reprograming the transcriptomes shifts from growth to defence. Overall, *ADR1s* emerge as key modulators in balancing plant immunity and growth.

### Chromatin Accessibility Dynamics and Transcriptional Reprogramming Orchestrate Immune Gene Activation and Growth Suppression in Plant Autoimmunity

To investigate chromatin modifications underlying transcriptional reprogramming during autoimmunity, we profiled chromatin accessibility in *DM1-DM2d*, *DM6-DM7*, and WT plants using Assay for Transposase-Accessible Chromatin using sequencing (ATAC-Seq). We observed expected enrichment of ATAC-Seq reads near transcription start sites (TSS), confirming reliable mapping (**Figure S9A**). PCA revealed distinct chromatin accessibility landscapes between *DM* lines and WT, with *DM1-DM2d* exhibiting more pronounced shifts along PC1 than *DM6-DM7* (**Figure 6A**). After identifying Open Chromatin Regions (OCRs), we defined Differentially Accessible Regions (DARs) as OCRs with |log2FC| > 1 in *DMs* versus WT. Volcano plots demonstrated that regions with reduced chromatin accessibility significantly outnumbered those with increased accessibility in both *DM* lines, with this chromatin condensation being more pronounced in *DM1-DM2d* (**Figure 6B** and **S8B**).

**Figure 6.**
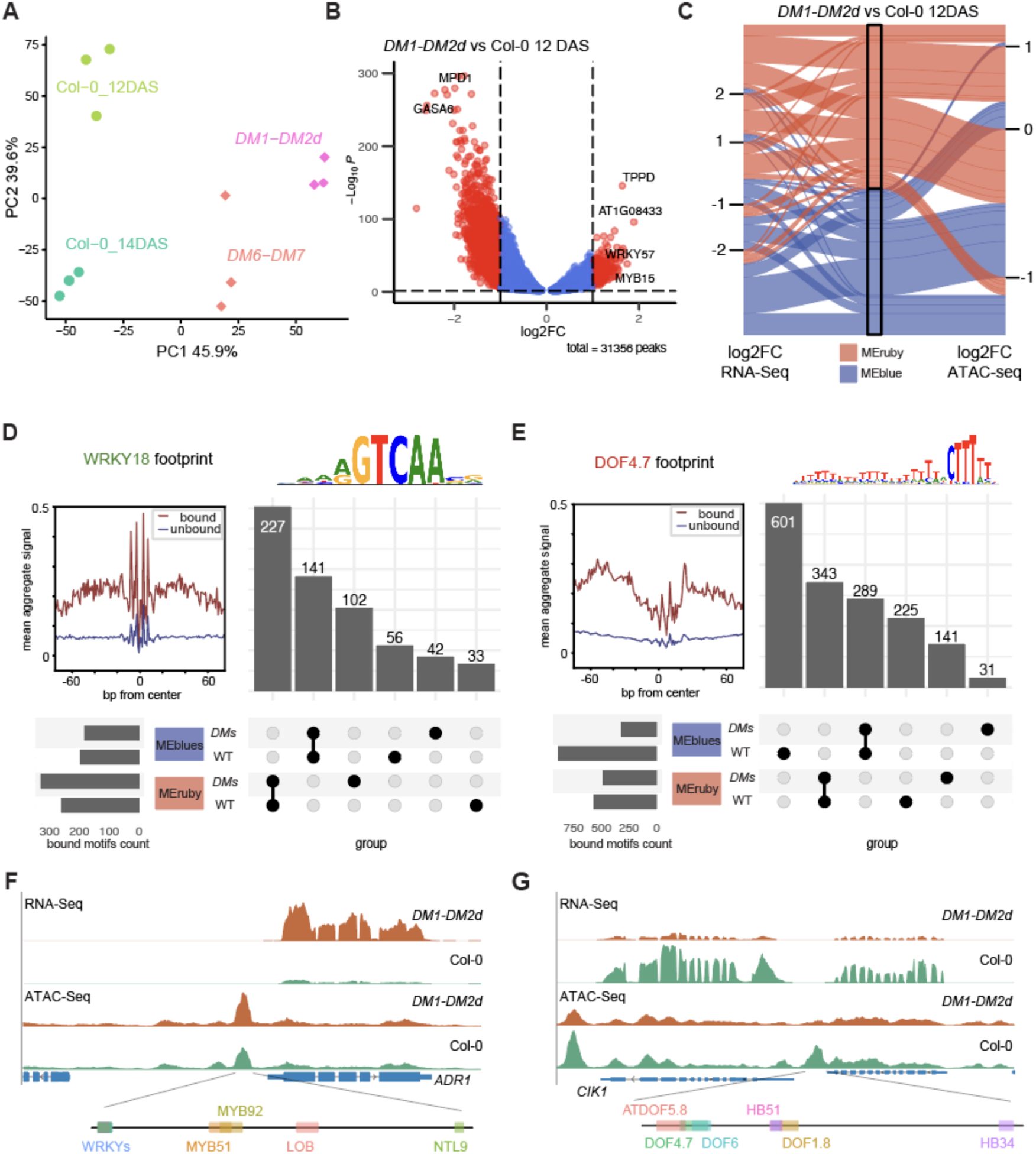
ATAC-Seq and RNA-Seq Co-analysis revealed TFs as activators and repressors in autoimmune transcriptome reprogramming. **(A)** Alluvia plot of correlation in RNA-Seq and ATAC-Seq. WGCNA modules MEblue and MEruby’s RNA-Seq and ATAC-Seq in *DM1-DM2d* vs Col-0 12DAS are plotted. **(B)** diffTF assigned TF into activator or repressor by the correlation of Differential TF activity and TF transcription level between WT Col-0 and *DMs*. TFBS, number of predicted TFBSs. ΔTF activity indicated the TFBS accessibility (x axis). p values are obtained through diffTF using the empirical approach and adjusted by the Benjamini-Hochberg procedure (y axis). **(C)** ChromVAR analysis of TFBS synergy and correlation. TFs are label as activators (green) or repressors (red). **(D)** Footprint and UpSet analysis of activator WRKY18, incorporated with predicted bound sites in WGCNA MEblues (MEblue + MEpaleblue) and MEruby modules across *DMs* and WT. **(E)** Footprint and UpSet analysis of repressor DOF4.7. **(F)** Visualization of RNA-Seq and ATAC-Seq in ADR1. ADR1 has a high transcription level in *DMs*, coincident with a more accessible ACR in its promoter, which contains multiple WRKY binding sites. **(G)** Visualization of RNA-Seq and ATAC-Seq in NIK3. Example of NIK3 contains repressor binding sites such DOF4.7. NIK3 is down-regulated in *DMs*, corresponding to a more closed chromatin. The reduced ACR in its promoter contains TFBS of repressor such as DOF4.7.

Integration of RNA-Seq and ATAC-Seq data using an alluvial diagram (**Figure 6C**) revealed contrasting patterns between growth and immune modules. Genes in the MEblue (growth-suppressed) exhibited widespread chromatin condensation, with 1638 of 1805 OCRs showing reduced accessibility (log2FC < 0). Conversely, the MEruby module (immune response) displayed a mixed pattern: fewer than half (777 of 1736) OCRs showed increased accessibility (log2FC > 0), whilst the majority (959 OCRs) exhibited decreased accessibility. GO enrichment analysis of DARs showed genes near upregulated DARs (log2FC > 1) were associated with systemic acquired resistance (SAR), whilst those near downregulated DARs (log2FC < −1) were linked to auxin response (**Figure S9C and S8D**). Growth-related genes undergo coordinated transcriptional repression and chromatin condensation, while immune-response genes show transcriptional activation despite variable chromatin accessibility changes. This suggests many defence gene loci may already exist in a permissive chromatin state, requiring only appropriate transcription factors to initiate expression (Luo et al., 2012; Ding and Wang, 2015; Ramirez-Prado et al., 2018).

TFs regulate gene expression by binding to specific DNA elements (Feng et al., 2016; Zou and Sun, 2023). In ATAC-Seq, TFs partially shield DNA from Tn5 transposase cleavage, creating subtle low-signal areas within accessible regions. We used TOBIAS for genome-wide detection of TF binding footprint (Bentsen et al., 2020) (**Figure S9E**). To understand TF behaviour in the context of chromatin dynamics during autoimmunity, we further used diffTF to integrate RNA-Seq and ATAC-Seq data and categorised TFs as either opener or closer based on the correlation between TF expression and chromatin accessibility at their predicted binding sites (Berest et al., 2019) (**Figure S9C**). This inference-based classification differs from the classical activator/repressor categorisation that relies on experimental assessment of transcriptional effects. When applying this integrated approach to *DM1-DM2d* and *DM6-DM7* autoimmunity contexts, we identified key TF families (**Figure S9F**, **Table S5**), including basic Leucine Zipper Domain (bZIP), TGACG SEQUENCE-SPECIFIC BINDING PROTEIN (TGA), DNA binding with one finger (DOF), WRKY, and NAM, ATAF1/2, CUC2 (NAC). Using an RNA-Seq |log2FC| > 2 threshold, we identified *WRKY18*, *TGA1* and NAC016 among highly up-regulated openers, while *DOF4.7*, *CDF5*, *COG1*, and *bZIP28* were the most up-regulated closers. ChromVAR assessed the interaction between TF-binding motifs in two dimensions: “correlation” (simultaneous accessibility of two motifs within an open chromatin region) and “synergy” (joint presence in accessible chromatin) (Schep et al., 2017), indicating minimal correlation and synergy between closers and openers (**Figure S9G**), suggesting mutual exclusiveness.

TOBIAS footprint analysis of *WRKY18*, a representative opener, showed steady binding patterns in MEblue-related modules across *DM* and WT samples. However, the MEruby module exhibited 102 additional binding sites in *DM* samples beyond the 227 sites shared with WT (**Figure 6D**). This increase aligns with *WRKY18*’s opener role, though the predominance of shared sites suggests these regions are already permissive without *DM* signalling. Conversely, *DOF4.7*, a closer, showed dramatically reduced binding (601 fewer sites) in MEblue in *DM* samples compared to WT, with only 289 sites maintained (**Figure 6E**). MEruby *DOF4.7* binding was characterised by 343 shared sites, with variable gains and losses in *DM* versus WT. These patterns are exemplified by two loci: the *ADR1* locus displayed an OCR near its TSS that was accessible in WT but became further opened in *DM1-DM2d*; conversely, the RLK *CLAVATA3 INSENSITIVE RECEPTOR KINASE 1* (*CIK1*) locus, which regulates stem cell homeostasis (Hu et al., 2018), showed pronounced closure at its *DOF4.7* motif containing proximal promoter in *DM1-DM2d* (**Figure 6F**). Similar patterns were observed across MEruby and MEblue modules (**Figure S10**).

These binding profiles suggest WRKYs and NACs activate immune gene expression through selective chromatin relaxation at key loci. Conversely, DOF4.7, COG1, and CDF5 facilitate chromatin condensation, suppressing growth-related genes. Paradoxically, despite elevated expression, these closers show reduced binding events in MEblue modules in *DM1-DM2d* versus WT (**Figure 6E**, **Figure S10B-D**), potentially because chromatin condensation at target sites limits TF accessibility.

### TF-Transcriptome Module Network Reveals Differential Mode of Action of Openers and Closers

To explore the regulatory interplay between individual TFs and transcriptional performance of their target genes, we constructed a TF-Transcriptome Module Network integrating RNA-Seq and ATAC-Seq data (**Figure 7A**). Candidate TFs playing critical roles in chromatin dynamics during immune responses were chosen by combining the DEGs obtained from RNA-Seq and diffTF results from ATAC-seq. This network links these TFs to changes in chromatin accessibility of their target motifs that belong to a certain WGCNA module, thereby connecting TF activities to representative WGCNA modules. The network diagram depicts a link between a TF as hexagonal node and a WGCNA module as diamond-shaped node while edges representing the enrichment of TF target sites in the connected WGCNA module. Node size and colour were used to represent expression changes of TFs in the *DM1-DM2d* line compared to WT, while edge colours indicate varying degrees of chromatin accessibility changes.

**Figure 7.**
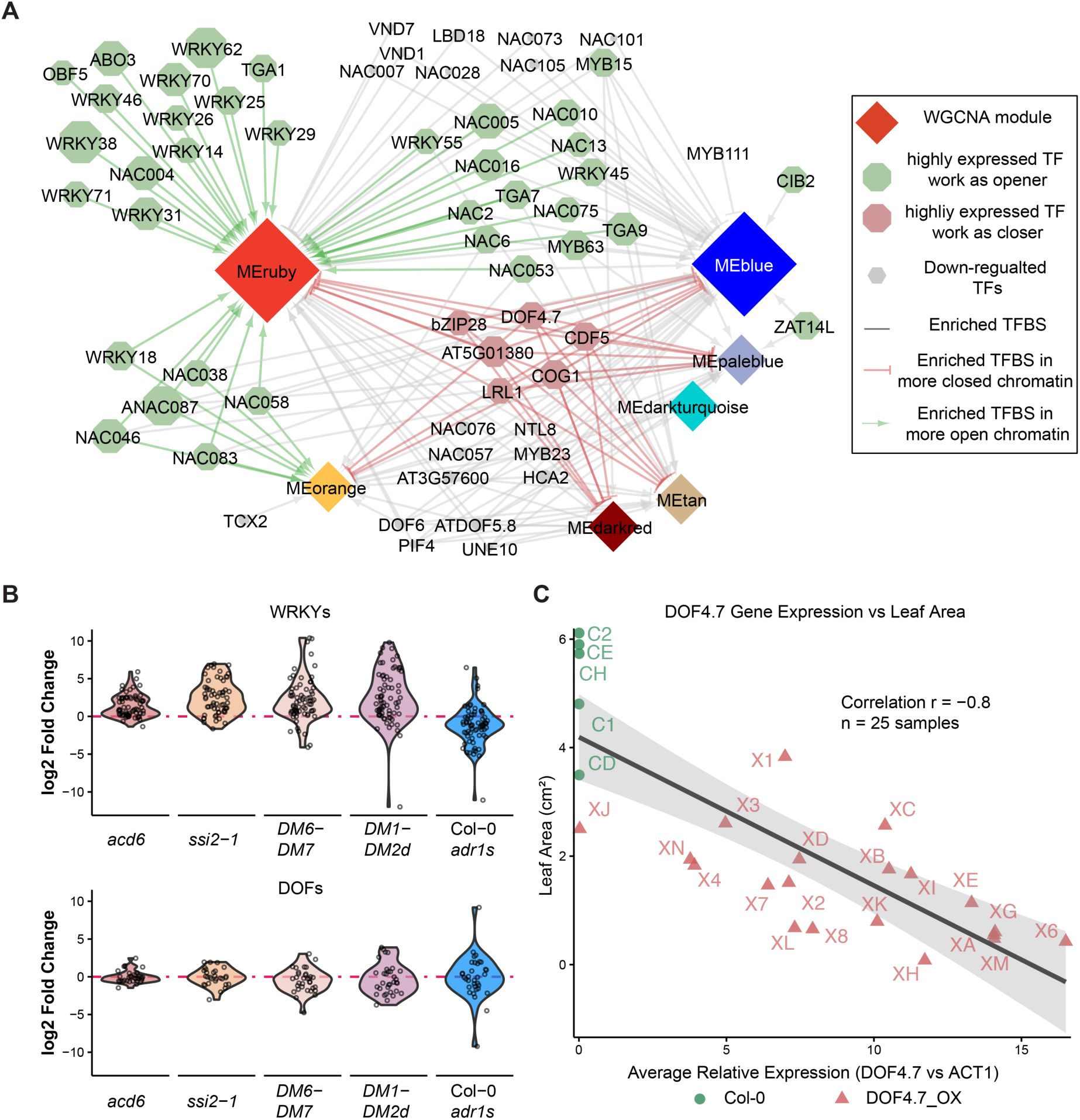
TF-WGCNA module Interaction Network and Model of *DMs* Immune Response. **(A)** TF-Module Network: This network diagram illustrates the complex interactions between TFs) and WGCNA modules identified in the *DMs* immune response. Diamond-shaped nodes represent WGCNA modules, with their size proportional to the number of genes they contain. Hexagonal nodes depict TFs classified as openers or closers by diffTF analysis. TF nodes are colored based on their expression changes in *DM1-DM2d* compared to WT: green hexagons represent upregulated openers (log2FC > 2), red hexagons indicate upregulated closers (log2FC > 2), and grey hexagons show openers or closers with less significant expression changes (log2FC < 2). Edges in the network represent enriched transcription factor binding sites (TFBS) for the connected TF within the respective module, predicted by Analysis of Motif Enrichment (AME) in MEME suite. The color of these edges reflects chromatin accessibility changes in *DM1-DM2d* compared to WT, as determined by ATAC-seq data. Green edges signify increased accessibility (more open chromatin) in *DM1-DM2d* vs WT, red edges indicate decreased accessibility (more closed chromatin), and grey edges represent no significant change in TF expression. **(B)** Violin plots show log₂FC distributions for WRKY and DOF. Expression changes are compared across four autoimmunity liens: *acd6*, *ssi2-1, DM6-DM7*, *DM1-DM2d* and Col-0 *adr1 triple*. Each violin shows the distribution density of log₂FC values, with individual data points overlaid as dots. **(C)** Functional validation of DOF4.7 as a growth suppressor. Scatter plot showing the relationship between *DOF4.7* transcriptional levels (relative to *ACT1*, x-axis) and plant size measured by PlantCV analysis (y-axis) across multiple independent overexpression T1 plants. The strong negative correlation (R = −0.8) demonstrates that elevated *DOF4.7* expression directly correlates with reduced plant size.

Our network analysis highlights a marked asymmetry in the regulation of immunity versus growth-related genes **(Figure 7A)**. Common immune responses (MEruby) are targeted by a multitude of upregulated opener TFs, chiefly WRKYs and NACs, with the MEorange module sharing several of these openers. Conversely, growth-associated processes (MEblues) are regulated by remarkably few closer TFs. Our initial analysis identified 121 openers and 51 closers (**Table S5**)—a notable numerical bias toward activators. This imbalance intensified in the network proper, where 34 openers but merely 6 closers met inclusion criteria of RNA-Seq up-regulation (**Table S6**). The consistent behaviour of WRKY family members as openers and DOF family members as closers was validated across multiple autoimmune systems (**Figure 7B**). Thus, despite their significantly smaller numbers, repressors appear to exercise disproportionate control over extensive portions of the repressed transcriptome during immune responses.

The number of opener TFs exclusively connected to MEruby outnumber the closer exclusively connected to MEblues. Most closers were connected to both modules, exerting widespread chromatin closing effects in both modules (**Figure 7A, red edges**). There were also a subset of opener TFs, including *NAC005*, *NAC006*, and *WRKY45*, which shows binding sites in both MEruby and MEblue modules. However, their activating influence was primarily exerted on MEruby, evidenced by increased chromatin accessibility (green edges) to MEruby but not MEblue (grey). This selective activation by shared openers as well as a large number of exclusive openers highlights an effective regulatory mechanism that is poised to simultaneously upregulate immune responses to amplify signalling. We note that functional redundancy among TFs classified as openers appears to be an important factor in amplifying immune signals. This redundancy is exemplified by WRKY members such as WRKY46 and WRKY70, which suppress JA-induced *PDF1.2* expression and enhance *P. syringae* resistance (Hu et al., 2012). Similarly, TGA1 and TGA4 promote SA biosynthesis, enabling full induction of *SARD1* and *CBP60g* in plant defence (Sun et al., 2018). In contrast, a small number of closer with large effects on both modules indicate an effective strategy to suppress growth-related processes. Local closure effects in the promoter elements of MEruby upon *DM* signalling also indicates sophisticated cross regulation between negative and positive promoter elements affecting transcription of a given gene. This notion is also supported by our discovery of some downregulated TFs, including some members of NACs and VNDs, like *NAC007* and *VND7*, maintaining strong connections to MEruby with closed status of their target sites (red edges at the interface with MEruby module). This means that local target sites of these TFs become inaccessible upon *DM* signalling concomitantly with their reduced expression, although their target genes are highly expressed. This suggests these TFs’ potential role in suppressing the target genes by occupying negative elements on the promoters in WT. Overall, the repressive links found in the network connected to both WGCNS modules suggest that a select group of TFs can effectively suppress numerous growth and development genes and at the same time facilitate transcription activation of defence gene by closing negative promoter elements, facilitating resource reallocation from growth to defence.

### *DOF4.7* Overexpression Confirms Growth-Specific Regulatory Mechanisms

To experimentally validate the predicted role of *DOF4.7* as a growth suppressor, we generated transgenic lines constitutively overexpressing *DOF4.7*. Multiple independent T1 transformants exhibited varying transgene expression levels, allowing us to assess the dose-dependent relationship between *DOF4.7* expression and plant growth (**Figure S13A-C**). Quantitative analysis revealed a strong negative correlation (R = −0.8) between *DOF4.7* transcript levels and plant size measured by PlantCV image analysis (**Figure 7C**). This robust relationship provides direct functional evidence that elevated *DOF4.7* expression constrains plant growth, confirming its classification as a growth suppressor. Importantly, *DOF4.7* overexpression did not significantly alter basal expression of immune regulators *ADR1* or *EDS16* (**Figure S13B,D**). The slightly altered *PR1* expression observed across overexpression lines (**Figure S13E**) reflects the inherent variability of this stress-responsive marker rather than immune activation.

The characterisation of *DOF4.7* as a growth suppressor with minimal impact on immune gene expression suggests that the growth-defence trade-off may be amenable to targeted manipulation through specific transcriptional regulators, offering potential strategies for crop improvement.

## Discussion

We utilised distinct *DMs* triggered autoimmunity to examine how NLR-mediated immunity disrupts growth-defence balance. By eliminating the helper *ADR1s* in *A. thaliana*, we identified their role in coordinating bifurcated signalling that both enhances immune responses and inhibits growth-related genes. Our analyses showed primarily contracted chromatin in growth-related genes, while immune-related genes displayed selective accessibility increases (**Figure 6B, Figure S6B**). This asymmetric chromatin remodelling pattern accompanied hierarchical NLR gene activation and extensive transcriptional reprogramming affecting immune activation and growth suppression.

### Common Feature of Plant Autoimmunity: Concurrent Upregulation of NLRs and TIR-containing Genes

Our *DM* systems provide a unique model where NLR activation triggers immune responses resembling ETI without PTI stimulation—comparable to effector-expression system Ngou et al., 2021 but achieved through incompatible gene pairs. Prolonged NLR signalling enables studying constitutive immune activation effects. Our meta-analysis revealed *DM*-triggered autoimmunity shares features with and differs from pathogen-induced responses. The MEorange module, uniquely upregulated in *DMs* but not pathogen-treated samples, contains LRR-RLPs, TIR-NBS proteins, and ion/calcium channels, reflecting physiological responses specific to chronic autoimmunity. Conversely, the MEdarkred module rich in JA-responsive genes showed activation in pathogen-treated samples but suppression in *DM* lines, highlighting fundamental differences between these immune contexts.

A key characteristic of *DM*-triggered autoimmunity is widespread NLR and TIR-encoding gene upregulation. TNL-associated *DMs* (*DM10-DM11* and *DM1-DM2d*) exhibited stronger TIR-containing gene induction than CNL/RPW8-associated *DM6-DM7*, correlating with autoimmune phenotype severity. This systematic TIR-NLR co-elevation indicates functional interdependence in immune signal amplification rather than random transcriptional dysregulation.

Our transcriptomic analysis connects with recent structural and biochemical findings that TIR domains function as NAD⁺ cleaving enzymes generating signalling molecules including nicotinamide and ADP-ribose variants (Horsefield et al., 2019). The differential ADR1s dependency between TNL-triggered (*DM1-DM2d*) and CNL-triggered (*DM6-DM7*) autoimmunity aligns with known activation mechanisms: TNLs typically require helper NLR-containing complexes like EDS1-PAD4-ADR1, while CNLs function more autonomously through direct resistosome formation (Förderer et al., 2022a; Huang et al., 2022; Jia et al., 2022; Jia et al., 2022). This mechanistic divergence appears particularly relevant given that TIR signalling actively boosts PTI (Tian et al., 2021). Our observed massive TIR-containing gene upregulation likely amplifies immune signalling through the EDS1-PAD4-ADR1.

### Distinct Immune Profiles in *DM*-triggered Autoimmunity versus Pathogen-induced Immunity

The *DM* specifically activated MEorange module represents a distinct autoimmune signature that reveals how immune activation becomes self-perpetuating through an interconnected network of cell surface surveillance, calcium signalling and transcriptional feedback loops. Central to this network is coordination between cell surface receptors and ion channels. Heightened expression of RLKs and RLPs, including *RLP34*, *RLP37*, *RLP47* and *SNC4*, suggests these surface-residing immune receptors establish a sustained surveillance system that perpetuates the autoimmune state (Zhang et al., 2014; Albert et al., 2015; Steidele and Stam, 2021). The concurrent enrichment of ion-permeable channels such as ACD6 and CNGC11 suggests that surface surveillance activates calcium flux to sustain NLR signalling (Lu et al., 2003b; Urquhart et al., 2011; Kim et al., 2022; Chen et al., 2023). Natural variation in *ACD6* dramatically influences immunity across *A. thaliana* accessions, acting as a key trade-off switch that can cause hybrid necrosis at low temperatures (Todesco et al., 2014; Świadek et al., 2017).

This immune network is further reinforced by intracellular components, including *ANK* and *DOWNY MILDEW RESISTANT 6* (*DMR6*), an oxygenase that regulates SA-dependent defence pathways (Zeilmaker et al., 2015). Cell wall remodelling enzymes, including *BGL2* (*PR2*), *TBL40*, *ECS1* and *PAE8*, further reshape cellular architecture to maintain the defensive state (Aufsatz et al., 1998; Bischoff et al., 2010; de Souza et al., 2014).

This self-reinforcing autoimmune network distinctly contrasts with pathogen-induced immunity. The MEdarkred module, enriched for JA responses, is uniquely upregulated during pathogen treatment but downregulated in *DM*s. During *P. syringae* infection, the bacterial toxin coronatine (COR) mimics JA-isoleucine, hijacking the *CORONATINE INSENSITIVE 1* (*COI1*) pathway (Zheng et al., 2012) and suppressing SA-dependent defences through JA-SA antagonism, as evidenced by JA-responsive genes including *PDF1.2*, *VSP1* and *VSP2* in this module (Brooks et al., 2005). Our flood assay, ensuring uniform pathogen exposure, clearly delineates pathogen-induced from autoimmune responses—contrasting with infiltration methods that often blur boundaries between ETI and localised acquired resistance (LAR) (Jacob et al., 2023).

### ADR1s as Master Regulators of Growth-Defence Balance

Our meta-analysis revealed ADR1s’ dual regulatory function in plant immunity. ADR1s maintain basal expression of immune-related genes (NLRs, RLPs, defence-related RLKs) whilst actively suppressing growth-related genes during immune activation. This suppressive role was evident when *adr1 triple* mutations reversed growth-related gene suppression in *DM* backgrounds (**Figure 3B-3C**).

The identification of robust transcriptome modules that capture the growth-defence trade-off across diverse immune contexts represents a significant advance in understanding plant immune regulation. The conservation of MEruby and MEblue module patterns across independent studies, spanning different pathogen treatments, autoimmune mutants, and genetic backgrounds, demonstrates that these modules reflect fundamental biological processes rather than experimental artefacts. Particularly noteworthy is the consistent linear relationship between growth suppression and immune activation (**Figure 3C, Figure S7B-C**), suggesting that plants operate along a defined regulatory axis. The ability of *adr1 triple* mutations to shift this balance towards growth whilst maintaining some immune capacity indicates that the trade-off, whilst robust, is not immutable and can be modulated through targeted genetic interventions.

Recent work by Chhillar et al., 2025 align with our results, demonstrating that the EDS1-PAD4-ADR1 node maintains immune component abundance and restricts pathogen growth, while the EDS1-SAG101-NRG1 node primarily regulates hypersensitive cell death (Jia et al., 2022; Chen et al., 2022). The necessity of knocking out all three ADR1 family members to fully reveal their collective impact underscores their overlapping yet crucial roles in immune signaling (Dong et al., 2016; Saile et al., 2020).

Our discovery that ADR1s simultaneously bolster immune readiness whilst restraining growth-related gene expression establishes a framework for understanding how plants navigate the growth-defence interplay. The dual regulatory role of ADR1s provides important molecular insights into this trade-off, traditionally thought to be resource-limited but now understood as modulated through various signalling mechanisms (He et al., 2022). Our analysis reveals a striking linear correlation between growth suppression and defence activation across diverse conditions, demonstrating that ADR1s coordinate these opposing processes with remarkable consistency. Given that regulatory motifs of defence and growth TFs downstream of ADR1 are largely non-overlapping (**Figure S9G**), these pathways exist as separate modules despite their coordinated regulation. By modifying the link between ADR1s and their downstream growth modules, it may be possible to engineer a plant that deviates from the cline (**Figure 3C**) to limit the growth cost of infection while maintaining potent ADR1-mediated immune responses.

### Interplay of Chromatin Accessibility and TF Activity in Immune Response

The analysis reveals an asymmetric transcriptional network: redundant opener TFs versus few closer TFs controlling vast repressed regions. One intriguing finding through ATAC-Seq analysis is that whilst MEblue growth module genes showed widespread chromatin condensation, not all MEruby immune module genes displayed increased accessibility in *DM* lines (**Figure 6**). Recent single-cell research (Nobori et al., 2025) corroborates our findings, showing that transcriptional activation of defence genes frequently occurs without detectable changes in chromatin accessibility regions. Many genes exist in a permissive chromatin state, requiring only appropriate transcription factor binding to initiate expression during immune responses (**Figure 6**).

Motif-based predictions and accessibility mapping offer insights into the immune regulatory landscape. Though binding site predictions require experimental validation, our TF-network analysis (**Figure 7A**) revealed patterns consistent with established immune regulatory mechanisms. The network showed enrichment of WRKY, NAC, and bZIP binding motifs—key families previously implicated in immune regulation (Tsuda and Somssich, 2015). While WRKY binding predictions may be overrepresented due to their short consensus motifs (W-box: TTGACC/T), their consistent enrichment across immune-responsive loci supports their biological relevance.

Conversely, MEblue module analysis identified a distinct class of potential regulators—DOF transcription factors (*CDF5*, *COG1*, *DOF4.7*) classified as “closers” in our diffTF analysis. While DOF family members can function as transcriptional repressors in specific contexts— CDF5 repressing CONSTANS expression in photoperiodic flowering (Fornara et al., 2009; Henriques et al., 2017), DOF4.7 negatively regulating floral organ abscission and rosette size (Wei et al., 2010), and COG1 acting as a negative regulator in phytochrome-mediated light responses (Park et al., 2003) — our analysis reveals a novel chromatin-based regulatory mechanism. Unlike other well-known EAR motif-containing TFs that recruit Polycomb Repressive Complex 2 (PRC2) for chromatin marking (Baile et al., 2021), these DOFs lack EAR domains yet correlate with widespread chromatin condensation at target loci during immune responses.

The “closer” classification of DOF transcription factors (*DOF4.7*, *COG1*, *CDF5*) reflects their paradoxical behaviour: despite being upregulated in *DM* lines, they correlate with reduced chromatin accessibility at their predicted binding sites. This apparent contradiction suggests that elevated DOF expression drives chromatin condensation at target promoters, effectively silencing growth-related genes. The shared binding motif (AAAG core) among these DOF factors enables coordinated targeting of growth-associated loci, with their collective upregulation creating a repressive chromatin landscape that broadly suppresses developmental processes during immune activation. The strong negative correlation (R = −0.8) between *DOF4.7* expression and plant size across multiple independent transgenic lines demonstrates that this transcription factor functions as a *bona fide* growth suppressor (**Figure 7C, Figure S13**). The specificity of this effect—evident from unchanged basal immune gene expression (*ADR1*, *EDS16, PR1*) in overexpression lines—confirms that DOF4.7 primarily affect growth-related pathways rather than broadly affecting cellular metabolism (**Figure 7C, Figure S13**).

The concentrated action of these few DOF “closers” contrasts sharply with the distributed network of numerous WRKY and NAC “openers”, revealing an asymmetric regulatory architecture that coordinates growth-defence transitions. The molecular mechanisms underlying DOF-mediated chromatin condensation require further investigation, though the correlation between DOF upregulation and widespread growth gene silencing suggests their central role in immune-triggered developmental suppression.

### Limits of this Study and Future Plan

Bulk tissue analysis reveals broad transcriptional landscapes, representing merely a snapshot of dynamic processes. Future single-cell approaches (Rich-Griffin et al., 2020) would illuminate how individual cell types orchestrate the growth-defence balance, whilst time-course analyses could delineate the regulatory cascade establishing autoimmunity. The computational network analyses, built upon predicted TF binding sites and module associations, require experimental validation of proposed mechanisms.

While DOF transcription factors (*DOF4.7*, *COG1*, and *CDF5*) emerged as potential growth suppressors with remarkable influence over growth-related loci, future investigations could explore whether targeted manipulation of these DOF factors might decouple growth suppression from immune activation. Collective knockout of these DOF transcription factors, alongside strategic manipulation of TIR-encoding genes or ion-channels across diverse genetic backgrounds, represents a promising approach to restore balance between growth and immunity.

---

**Figure S1.**
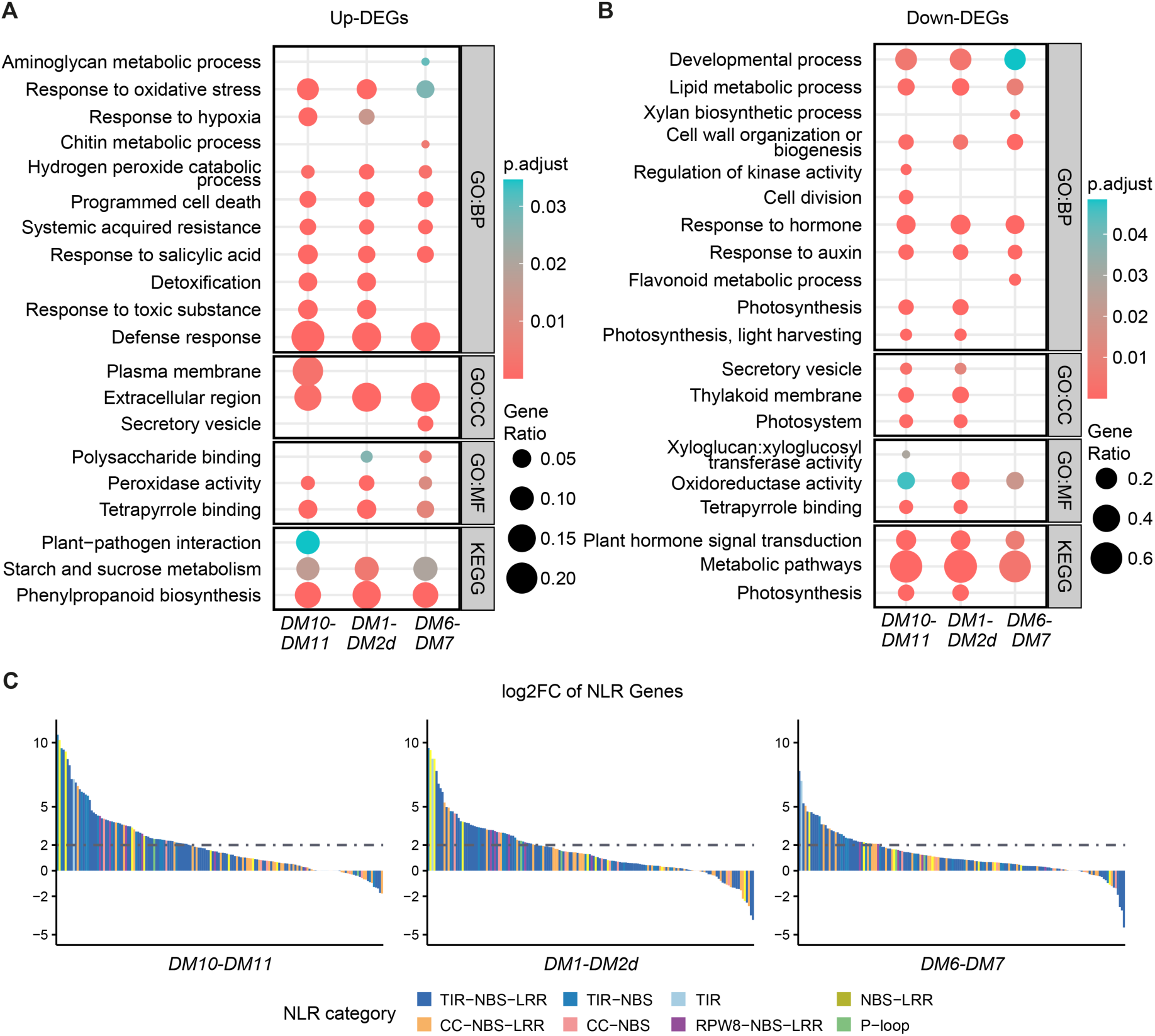
The GO terms and NLR profile. **(A)** The Gene Ontology (GO) terms of top up-regulated genes (up-DEGs) in three *DMs*. **(B)** The GO terms of top down-regulated genes (down-DEGs) in three autoimmune *DMs*. **(C)** Different expressions of NLRs between autoimmune lines and their WT control, with log2FC > 2, adjusted p-value < 0.05, baseMean > 10 as cutoff, among 167 NLRs, *DM10-DM11* had 59 NLRs highly up-regulated, *DM1-DM2d* had 46 NLRs highly up-regulated, *DM6-DM7* had 34 NLRs highly up-regulated. NLRs list was adopted from (Guo et al., 2011).

**Figure S2.**
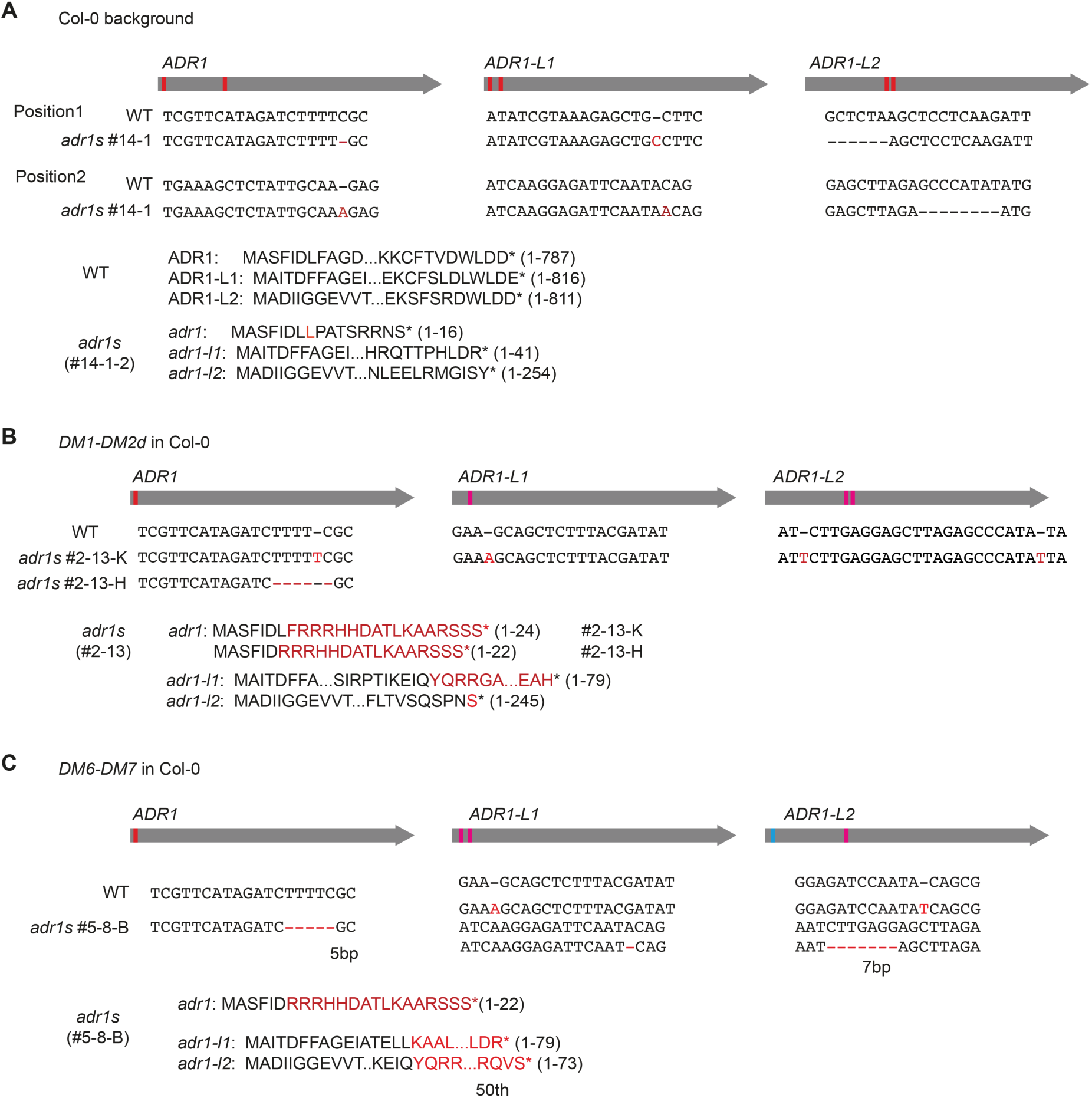
Illustration of mutations of *adr1s triple* in different backgrounds. **(A)** Col-0 background. **(B)** *DM1-DM2d* in background. **(C)** *DM6-DM7* in Col-0 background.

**Figure S3.**
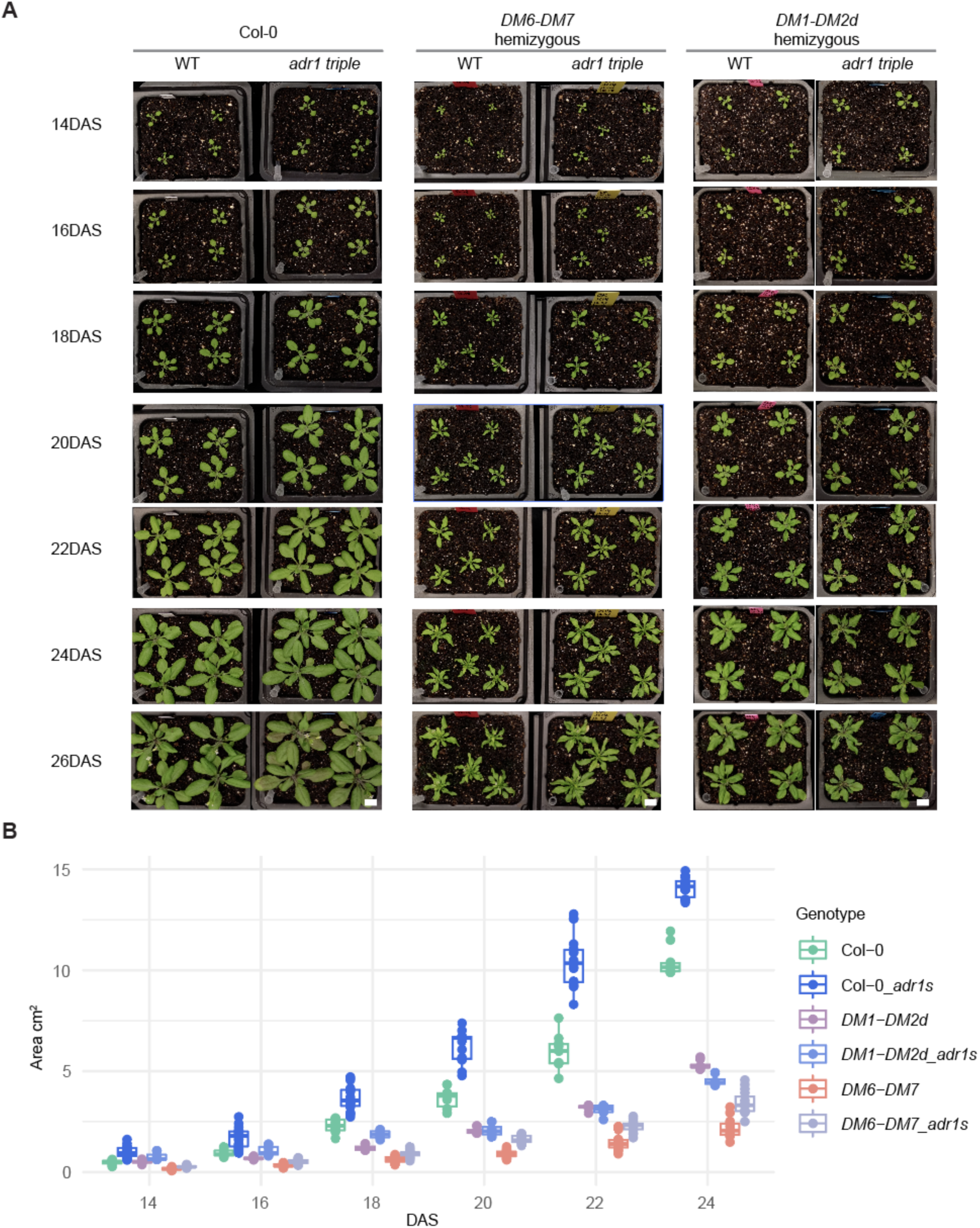
Knock-out of ADR1s increased plant size and alleviated autoimmunity phenotype in *DM6-DM7* and *DM1-DM2d*. **(A)** Growth of WT and *adr1 triple* in soil. The seeds were germinated on ½ MS plates supplemented with 1% sucrose and grew till 6 DAS before being moved to the soil. **(B)** Quantification of plant size. Plant size was quantified by PlantCV using photos.

**Figure S4.**
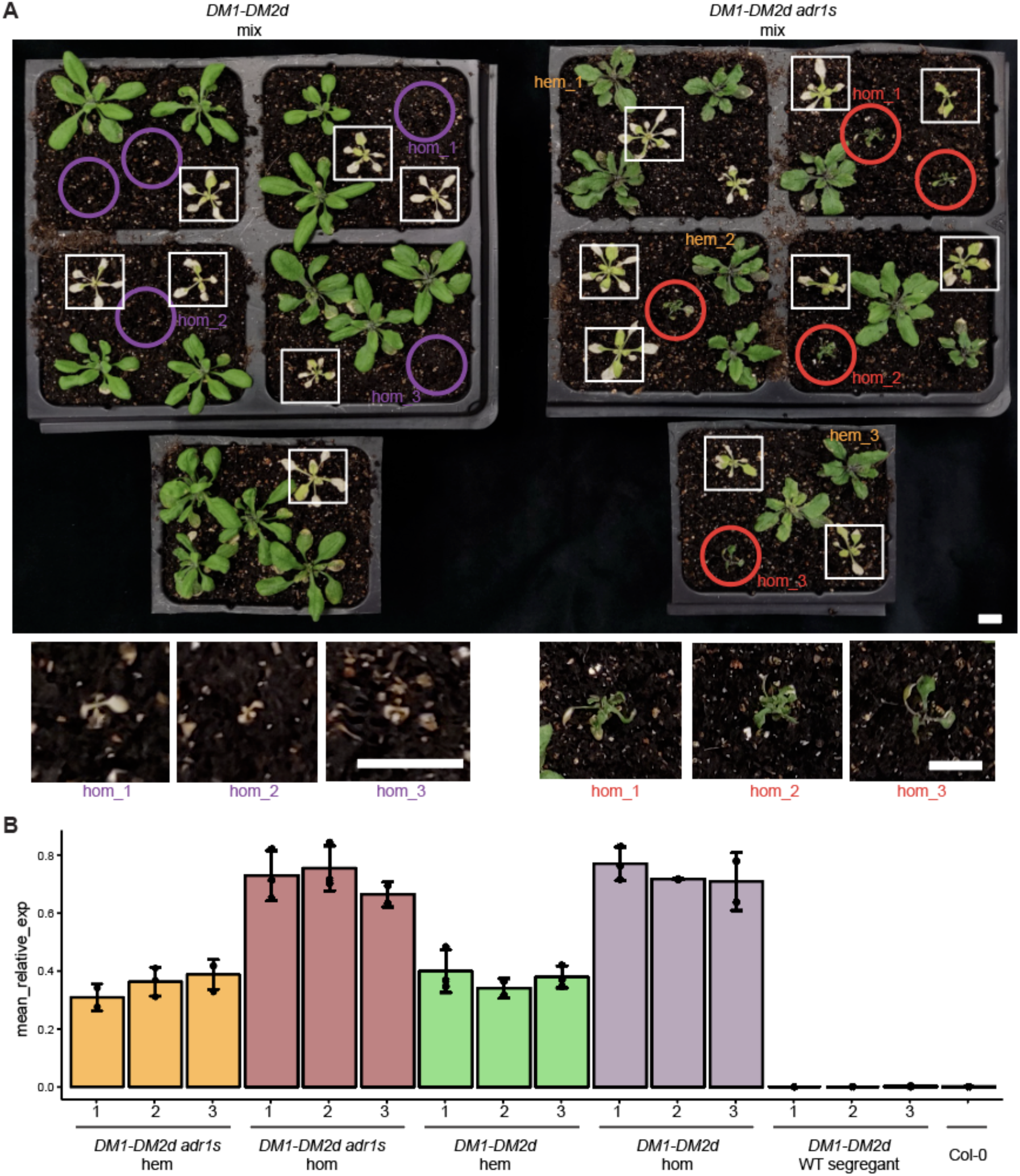
Phenotypic comparison and copy Number Variation (CNV) analysis of *DM1-DM2d* and its *adr1 triple* Mutant. **(A)** Phenotypic differences between *DM1-DM2d* and *DM1-DM2d adr1 triple* mutant plants. Homozygous *DM1-DM2d* plants displayed severe autoimmune phenotypes, including chlorosis and stunted growth at early stage, compared to milder symptoms in hemizygous plants. Partial rescue of these phenotypes was observed in *DM1-DM2d adr1 triple* mutants, though homozygous plants still exhibited more pronounced symptoms than their hemizygous counterparts. Circles indicate homozygous plants, which are shown in zoomed-in images in the lower panel. The white rectangles mark withered WT-segregants after spraying Basta (as *pDH666* carries the Basta resistance gene). Scale bars represent 5 mm. **(B)** Transgene copy number analysis using qPCR of *pDH666* (*DM1-DM2d* construct) relative to the reference gene EDS16, which is present in two copies in diploid *A. thaliana*. Homozygous plants showed approximately twice the construct CNV compared to hemizygous plants, correlating with the severity of phenotypes. Bars represent mean relative CNV ± SD from three biological replicates.

**Figure S5.**
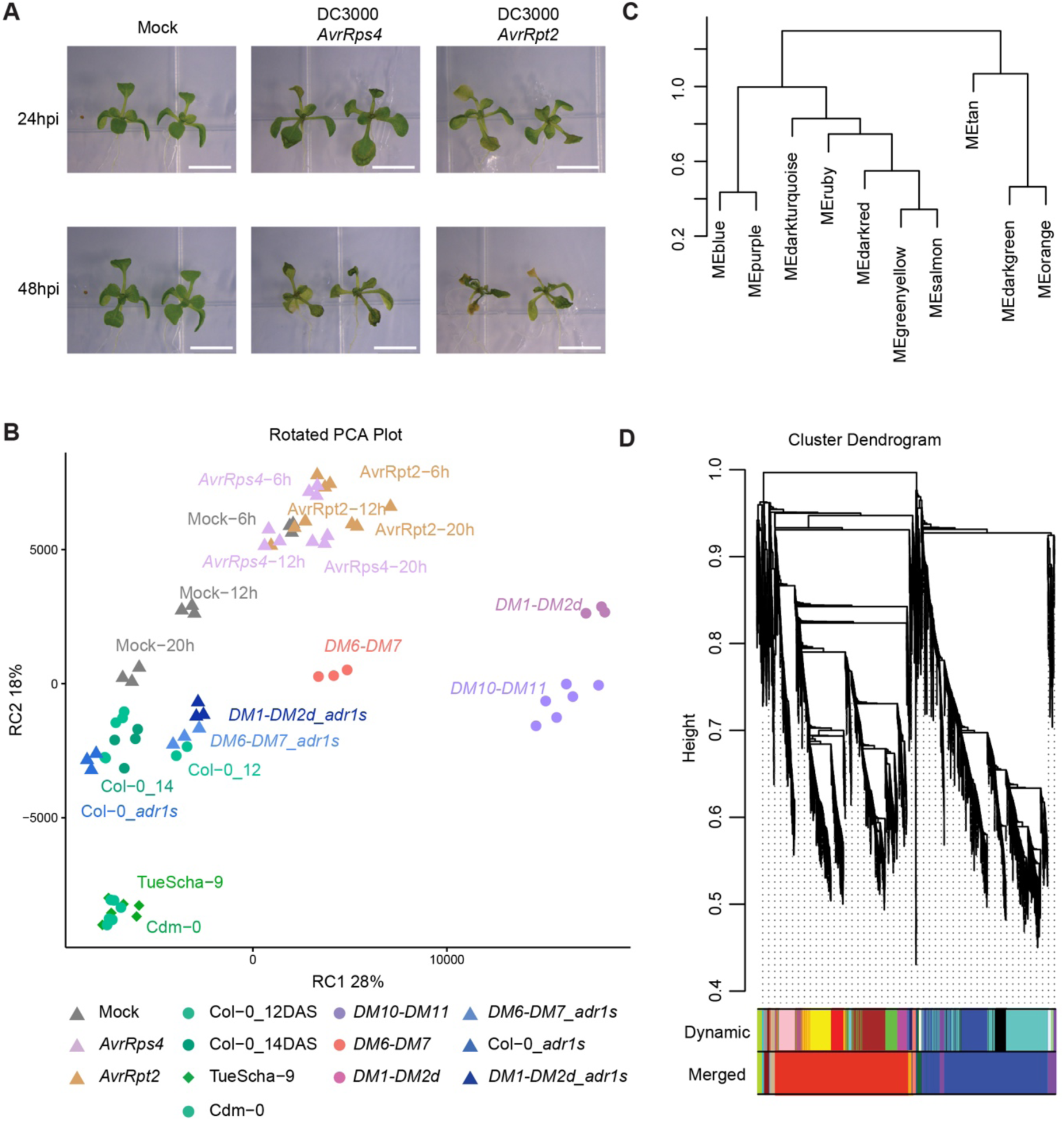
Co-expression analysis by WGCNA. **(A)** Phenotypes of plants following pathogen treatment by transient flood of solutions of DC3000 carrying *AvrRsp4* or *AvrRpt2*. The photos capture the appearance of treated plants at two time points: 24 hpi and 48 hpi. **(B)** Rotated Principal Component Analysis (Rotated PCA) of all samples used in the meta-analysis. The Rotated PC1 (RC1) corresponds to immune activation. **(C)** The dendrogram shows the hierarchical clustering of module eigengenes, identified by their corresponding colours. **(D)** The cluster dendrogram. The top 20% most variable genes (5454 out of 27270 genes) across 71 samples were identified using the median absolute deviation (MAD) metric and then analysed via the WGCNA. Variable genes were divided into ten modules after the dynamic tree-cut method and merging.

**Figure S6.**
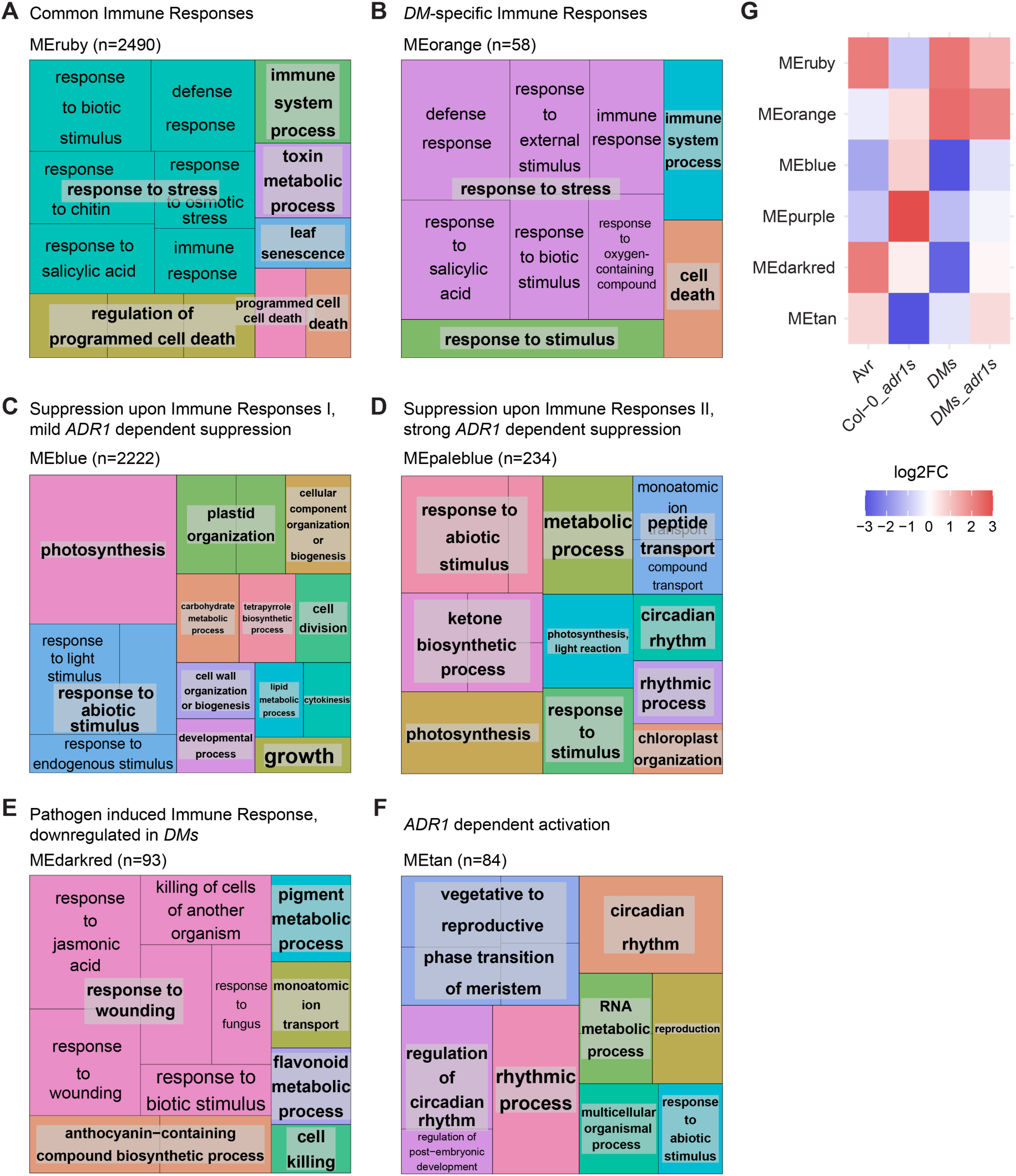
Treemap of enriched GO terms in WGCNA modules. **(A)** Common immune responses MEruby module and *DM*-specific immune responses MEorange module. **(B)** Suppression upon Immune Responses I, mild ADR1 dependent suppression MEblue and Suppression upon Immune Responses II, strong ADR1 dependent suppression MEpurple. **(C)** Pathogen induced Immune Response, downregulated in DMs MEdarkred and ADR1 dependent activation MEtan. The GO analysis is performed by AgriGO SEA and plotted with Revigo Treemap with customized R scripts. **(D)** the simplified modules heatmap.

**Figure S7.**
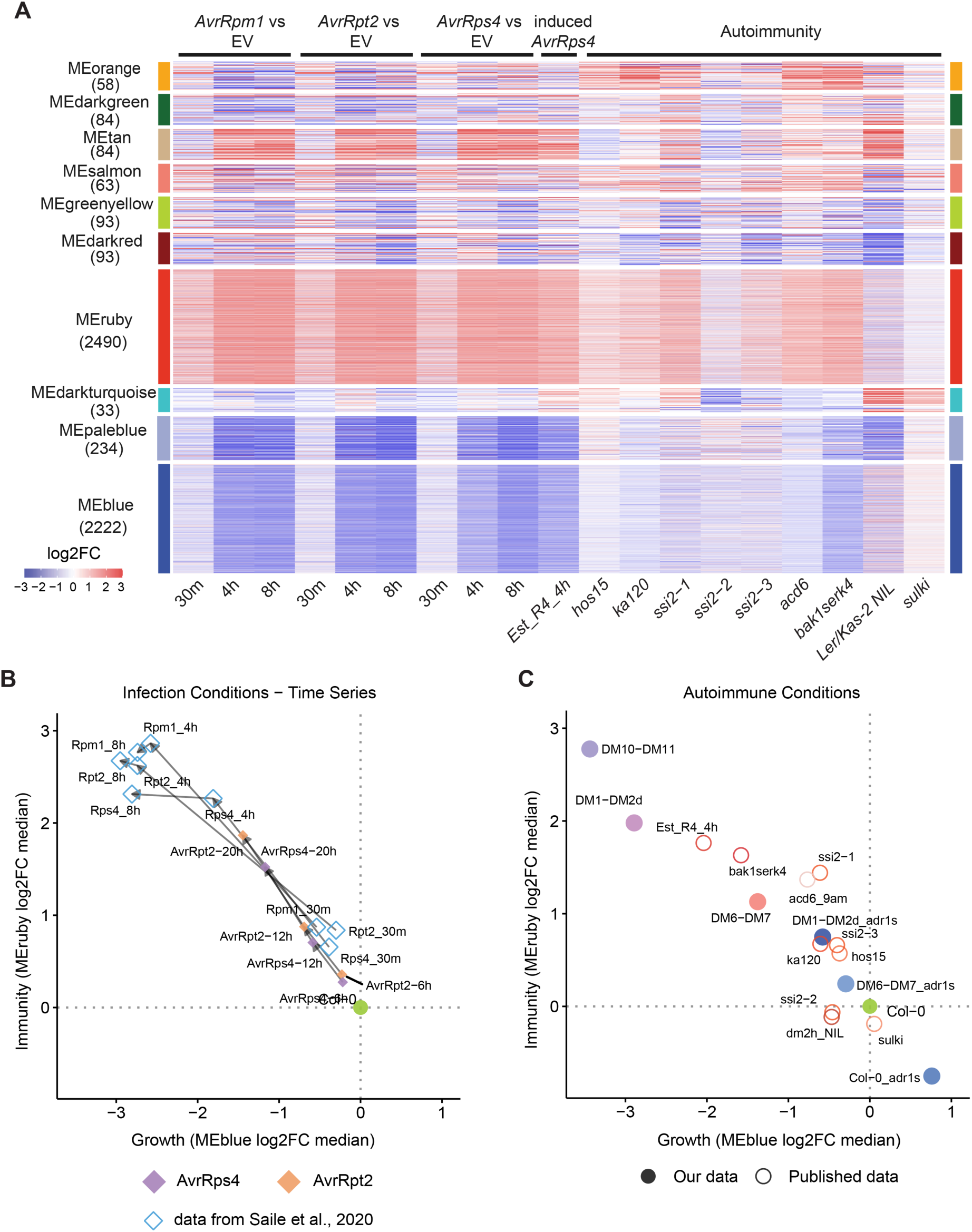
Validation of WGCNA modules across diverse transcriptome datasets highlights conserved immune response signatures. **(A)** Heatmap showing the behaviour of the WGCNA modules (initially identified in Figure 3) across multiple published transcriptome datasets related to plant immunity. These modules, derived from transient bacterial effector and NLR autoimmunity experiments (shown in Figure 3), were analysed here across independent datasets to validate their biological significance. The y-axis shows the same WGCNA modules (ME = Module Eigengene) as in Figure 3. The x-axis represents different experimental conditions from various studies including: time-course studies of bacterial effector responses (*AvrRpm1*, *AvrRpt2*, and *AvrRps4* vs *EV* by infiltration), estradiol-inducible *AvrRps4* system, and several autoimmune mutants. Colores represent log2FC in expression, with red indicating upregulation and blue indicating downregulation. This cross-validation across multiple experimental systems and genetic backgrounds confirms that these WGCNA modules capture fundamental and reproducible aspects of plant immune responses. Data were obtained from multiple published studies: bacterial effector responses (Saile et al., 2020), estradiol-inducible *AvrRps4* (Ngou et al., 2021), autoimmune mutants including *acd6-1* (Zhang et al., 2019), *bak1-4 serk4-1* (de Oliveira et al., 2016), *ssi2-1*, *ssi2-2* and *ssi2-3* (Yang et al., 2016), *hos15-4* (Yang et al., 2020), and genetic incompatibility studies comparing Ler/Kas-2 NIL with Kas-2 and its suppressor *sulki1-8* (Alcázar et al., 2009; Atanasov et al., 2018). **(B)** Scatter plot showing the reciprocal relationship between immunity activation (MEruby log₂FC median, y-axis) and growth suppression (MEblue log₂FC median, x-axis) in pathogen treatment time courses. Data points represent different pathogen treatments from our flood assay (*AvrRpt2* and *AvrRps4*, brown and purple) and Saile et al. infiltration experiments (blue circles) at various timepoints. Col-0 is positioned at the origin (0,0). The negative correlation demonstrates consistent trade-off dynamics irrespective of infection methodology. **(C)** Scatter plot illustrating the same growth-immunity trade-off across autoimmune backgrounds. Points represent our *DM* lines with and without *adr1 triple* mutations and various published autoimmune mutants including *acd6-1*, *bak1-4 serk4-1*, *ssi2* alleles, *ka120*, *hos15*, and Ler/Kas-2 NIL with its suppressor *sulki1-8*. The *adr1 triple* mutation shifts autoimmune transcriptomes toward baseline along the regulatory axis, confirming ADR1s as central modulators of this antagonistic relationship.

**Figure S8.**
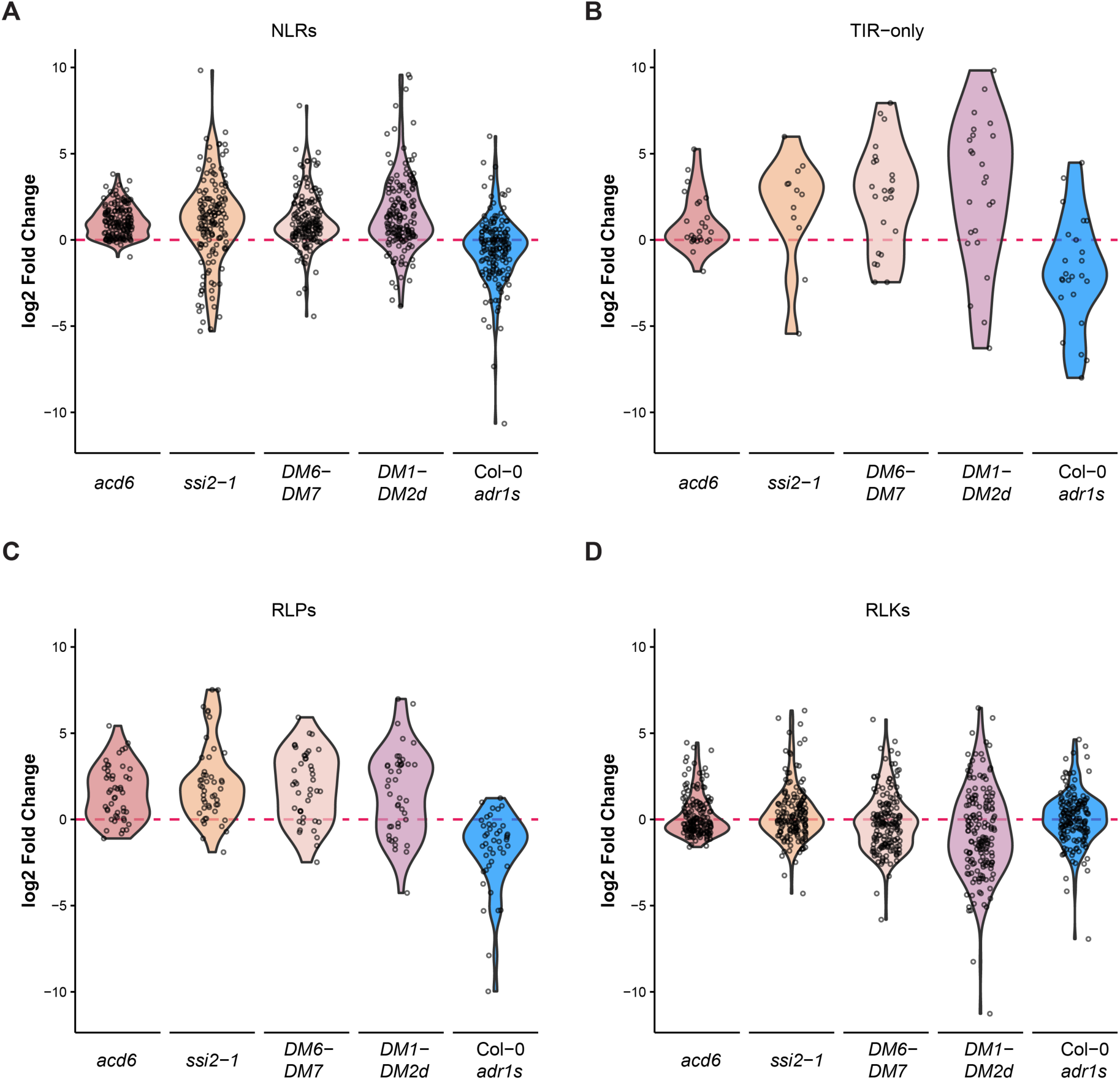
Differential expression patterns of key gene families across autoimmune mutants and *DM* lines. Violin plots show log₂FC distributions for **(A)** NLRs (nucleotide-binding leucine-rich repeat proteins), **(B)** TIR-only proteins (containing TIR domains without NBS-LRR), **(C)** RLPs (receptor-like proteins), and RLKs (receptor-like kinases). Expression changes are compared across five autoimmune lines: *acd6*, *ssi2-1*, *DM6-DM7*, *DM1-DM2d*, and Col-0 *adr1s* triple. Each violin shows the distribution density of log₂FC values, with individual data points overlaid as dots.

**Figure S9.**
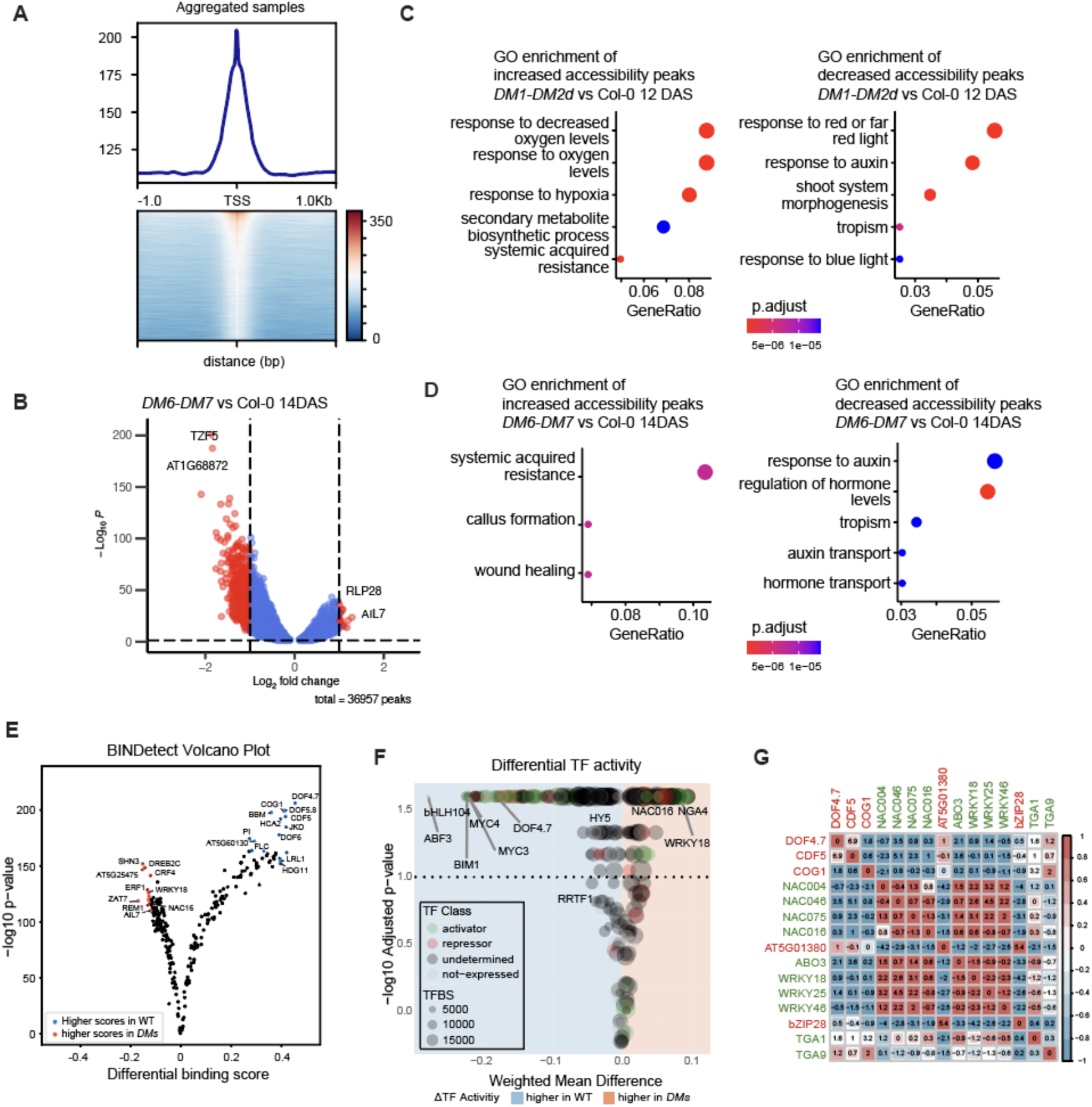
Analysis of ATAC-Seq Data in DM Autoimmunity. **(A)** Aggregated TSS enrichment plot for all ATAC-seq samples, illustrating chromatin accessibility near transcription start sites (TSS). The upper panel displays the aggregated signal, while the heatmap below indicates signal intensity around TSS regions, confirming data quality and accessibility mapping. **(B)** Volcano plot depicting differentially accessible chromatin peaks in DM6-DM7 versus Col-0 at 14 DAS. The x-axis represents log2FC of accessibility, and the y-axis shows the −log10 p-value. Highlighted peaks indicate significant changes in accessibility, with notable loci labeled. **(C)** Gene Ontology (GO) enrichment analysis for genomic regions with increased chromatin accessibility. The x-axis indicates GeneRatio, and the color gradient represents adjusted p-values. **(D)** GO terms enriched in regions with reduced chromatin accessibility. The x-axis represents GeneRatio, with the color gradient showing adjusted p-values. **(E)** Differential binding score analysis for transcription factor (TF) binding events using TOBIAS BINDetect. The x-axis displays the differential binding score, and the y-axis represents −log10 p-value. Key TFs with significant differences in binding activities between WT and DMs are highlighted. **(F)** Volcano plot integrating RNA-seq and ATAC-seq data. Weighted mean differences in TF activities (ΔTF activity) between WT and DMs are plotted on the x-axis, with −log10 adjusted p-values on the y-axis. TFs are categorized by their class (activator or repressor) and significance. **(G)** The heatmap represents TF binding motif correlation (color gradient) and synergy (numerical values in cells) across different transcription factors. Correlation values indicate co-accessibility of binding motifs, while synergy values quantify the co-occurrence of motifs within accessible chromatin regions. Key TFs are selected from diffTF result to their interactions in the DM and WT.

**Figure S10.**
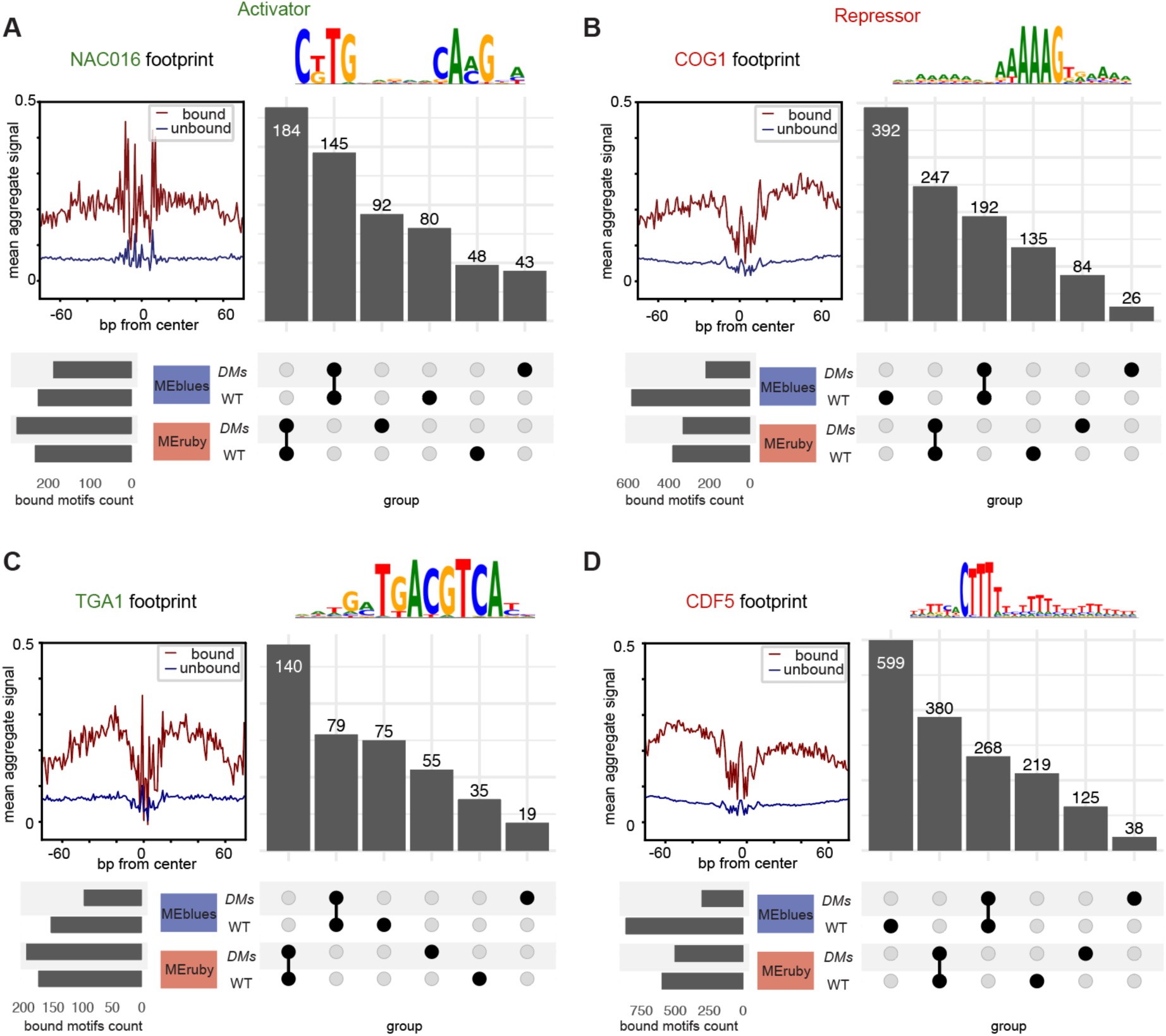
Footprint and UpSet analysis of activators and repressor. incorporated with predicted bound sites in MEblues (MEblue + MEpaleblue) and MEruby modules across *DMs* and WT. **(A)** NAC016 **(B)** COG1 **(C)** TGA1 **(D)** CDF5.

**Figure S11.**
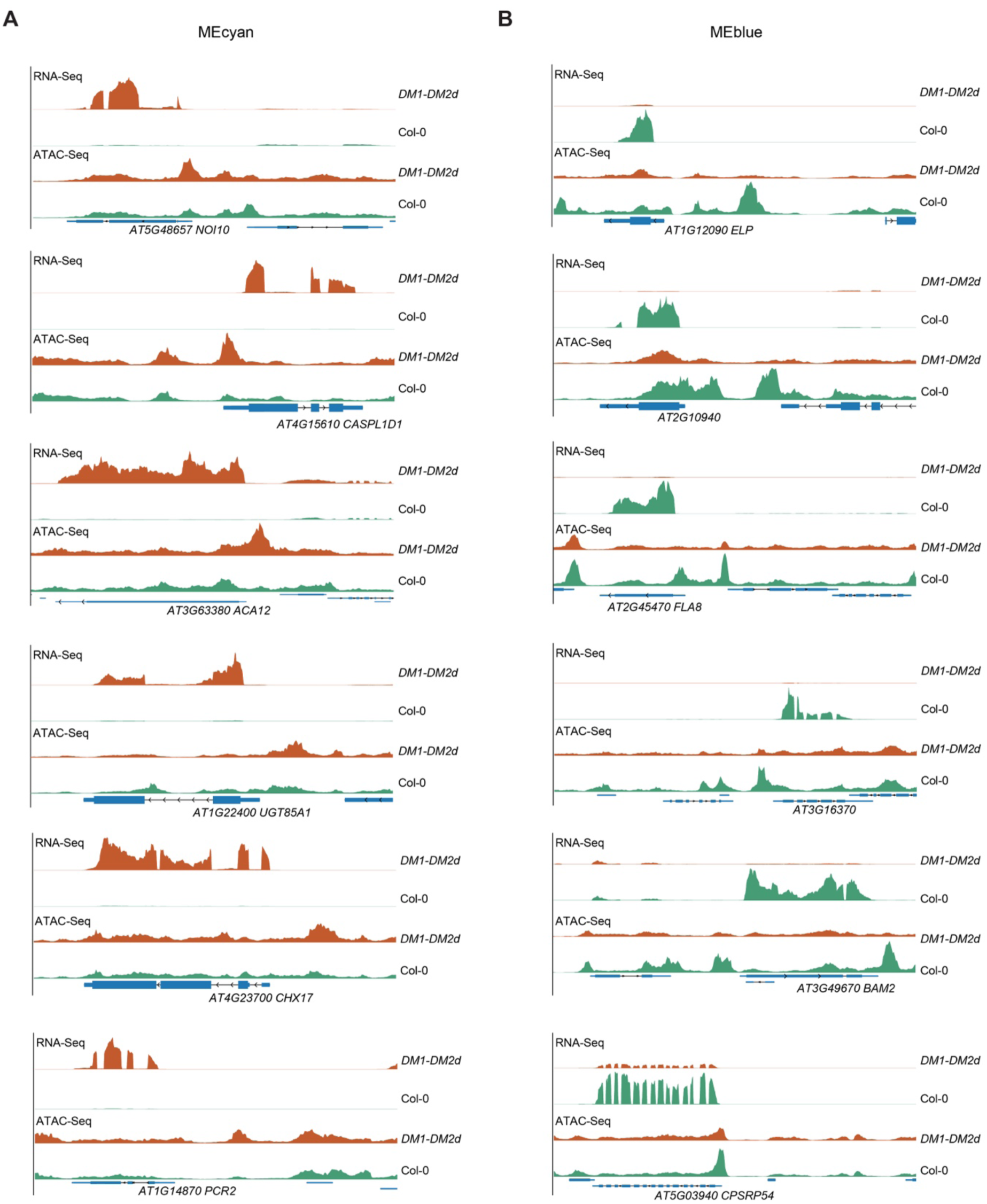
Example of genes from WGCNA modules. **(A)** Example of MEcyan up-regulated genes showing higher accessibility in ATAC-Seq. **(B)** Examples of MEblue down-regulated genes showing lower accessibility in ATAC-Seq.

**Figure S12.**
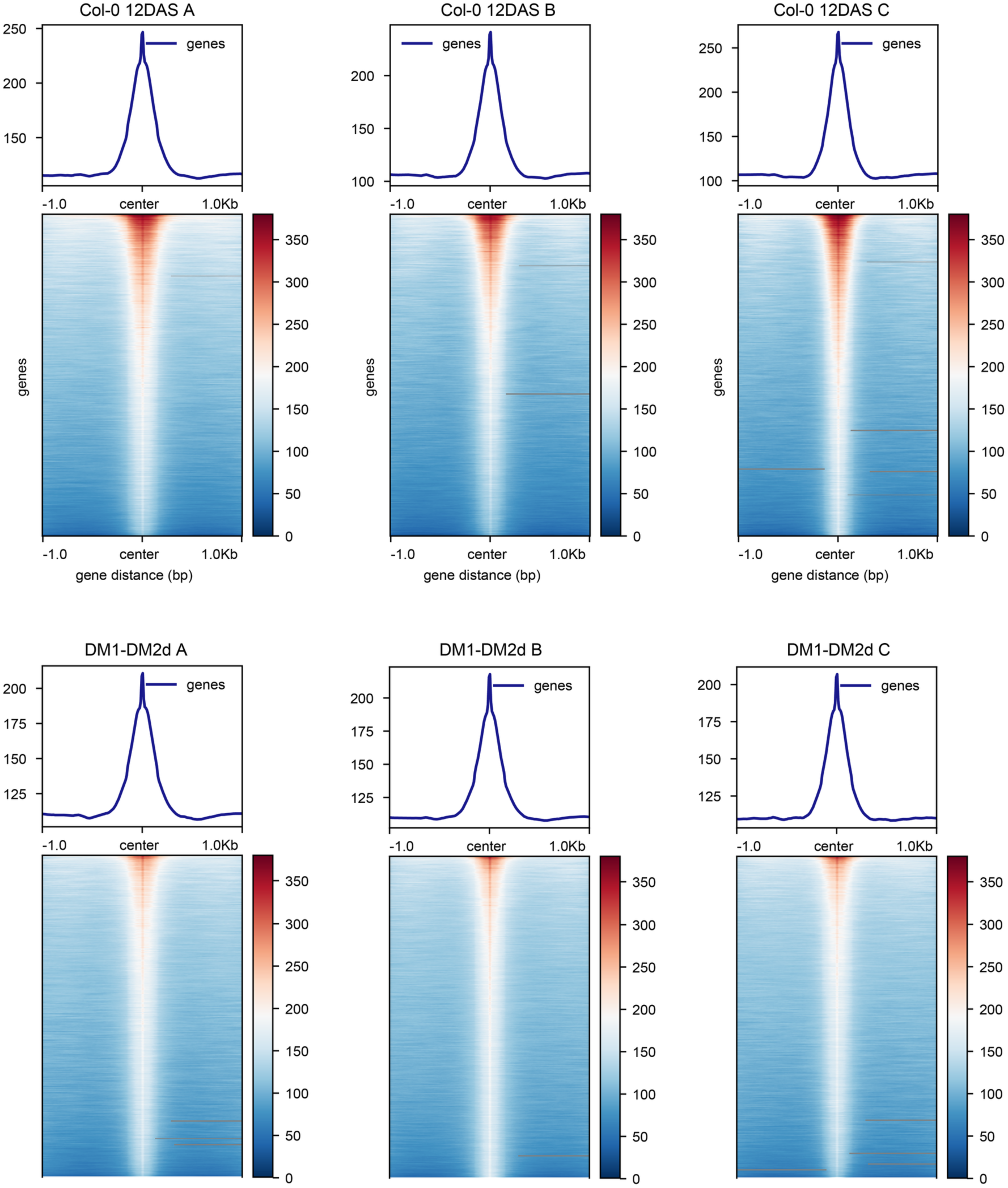
Quality control of ATAC-Seq.

**Figure S13.**
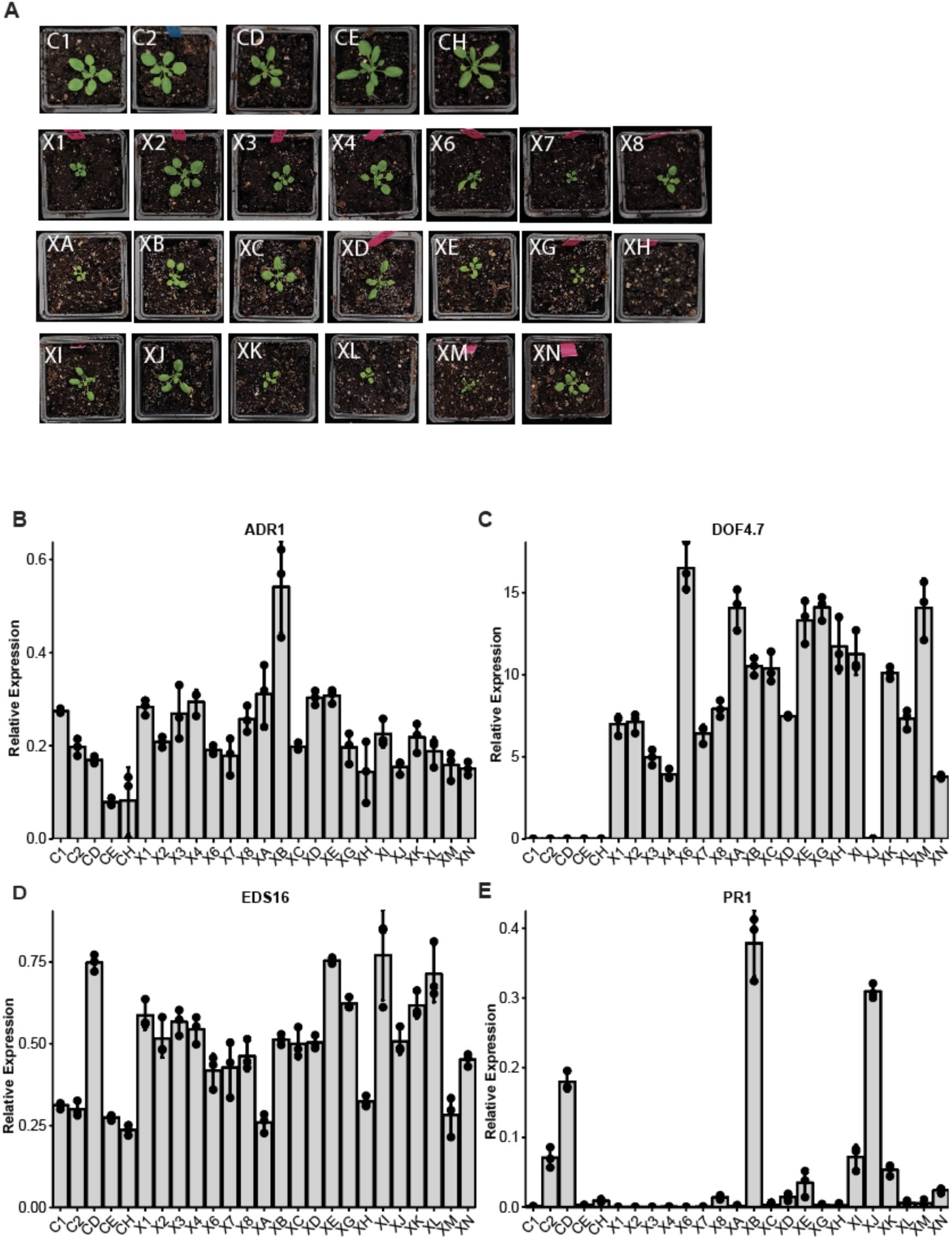
Phenotypic and molecular characterization of *DOF4.7* overexpression lines. **(A)** Representative photographs of *DOF4.7* overexpression T1 plants (DOF4.7 OX, naming start with “X”) lines and Col-0 controls (naming start with “C”) at equivalent developmental stages. Plant images are arranged in rows showing independent T1 transformants. Scale bars represent 2 cm. **(B-E)** Quantitative PCR analysis of gene expression in *DOF4.7* overexpression lines compared to Col-0 controls. Expression levels are normalised to the reference gene *ACT1*, with three technical replicates. Error bars represent standard deviation from three technical replicates. **(B)** *ADR1* **(C)** *DOF4.7*, same data used in Figure 7B. **(D)** *EDS16* **(E)** *PR1*.

## Methods

### Generation of *DM1-DM2d* lines and *DM6-DM7* lines

DM1 and DM2d with their native promoter were cloned into one construct DH666 by T5 Exonuclease-Dependent Assembly (TEDA) (Xia et al., 2019) and transformed into Col-0 by floral dip (Clough and Bent, 1998), mediated by *Agrobacterium* GV3101. The T1 seedlings were selected by Basta resistance. DM6-DM7 line was generated by transforming pWZ189 (DM6-DM7) into Col-0 and selected by Basta resistance. (Jinge Wang’s work)

*Hemizygous plants were used for generating seeds, given that the homozygous transformants cannot progress to the floral stage, thus resulting in a* Mendelian ratio of *WT-segregant: hemizygous plants: homozygous plants = 1:2:1 segregation*.

### RNA extraction and sequencing

The zygosity of *DM1-DM2d* seedlings on 12DAG, and the *DM6-DM7* seedling on 14DAG can be told by phenotype. The shoot/root of homozygous autoimmune plants or Col-0 were carefully cut by a blade, separately collected, frozen in liquid nitrogen. The samples were grounded and homogenised by TissueLyser II (QIAGEN). The total RNA was then extracted following LogSpin method (Yaffe et al., 2012). To confirm the homozygous plants in the p*ADR1s*-KO *DM1-DM2d* background, quantitative PCR (qPCR) was performed to detect any copy number variations using gDNA.

The extracted RNA was sent to Novogene or Azenta for quality control, strand-specific library preparation, and paired-end sequencing. Typically, 30M PE reads were generated for each *Arabidopsis* sample.

### mRNA-Seq Analysis

The genome TAIR10 and gene annotation from Ensembl were manually modified to add the insert sequence (pWZ189 and pHD666) to the end of files. Reads were mapped against modified genome using Hisat2 (Kim et al., 2019). Samtools (Danecek et al., 2021) was used to sort the reads, featureCounts (Liao et al., 2014) was used to assign reads to genomic features. The differentially expression analysis was performed by DEseq2 (Love et al., 2014).

### Transient pathogen flood assay

This protocol is modified from Ishiga et al., 2011. The procedure for the growth and inoculation of Arabidopsis seedlings and pathogen preparation and inoculation begins with the sterilization and stratification of Arabidopsis seeds. These seeds are then sown on 1/2 MS square agar plates and grown vertically in a plant tissue culture Percival at 22°C under long-day conditions. Six days post-sowing, normal-sized seedlings are transferred to new round plates of 1/2 MS for continued growth until the twelfth day after sowing, maintaining uniform inoculation. At this stage, the plates are placed horizontally. It is crucial to perform all these steps under sterile conditions. For the pathogen preparation, fresh pathogen is scraped from the plates 24-48 hours post-inoculation using pipette tips and resuspended in distilled water containing 10 mM MgCl2. The OD_600_ is adjusted to 0.1 for testing Avr triggered ETI. Silwet L-77 is added to this preparation just before use to achieve a final concentration of 0.028% (w/v). The pathogen inoculation involves the gentle pouring of the pathogen suspension onto plates to fully immerse the seedlings. After a brief immersion period of 2-3 minutes, the liquid is carefully drained off. The plates are then covered with lids and incubated at 22°C in a Percival for observation.

### ATAC-Seq

12 DAS DM1-DM2d and Col-0, 14 DAS DM6-DM7 and Col-0 shoots were used for ATAC-Seq. Nuclei were isolated from 0.05g LN snap-frozen plant tissues utilizing methods from Giuliano et al., 1988 and Dorrity et al., 2021. Tissues were pulverized at 30Hz for 40s with three 3mm beads in a TissueLyser II (Qiagen) within a 2mL EP tube, with subsequent procedures conducted on ice and centrifugation at 4°C. Buffer A (1.5mL; 0.8 M sucrose, 10 mM MgCl_2_, 25 mM Tris-HCl pH 8.0, 1x Protease Inhibitor) was quickly added to the powdered tissue, then gently mixed. After filtration through 40μm and 10μm strainers, the mixture was centrifuged at 2000 × g for 5 minutes. The resulting pellet was resuspended in 1mL Buffer B (0.4 M sucrose, 10 mM MgCl2, 25 mM Tris-HCl pH 8.0, 1x Protease Inhibitor, 1% Triton X-100), then loaded over a 25/75 Percoll gradient. The Percoll gradient is prepared by mixing Percoll with Buffer B. Following a 2500 × g, 15 min centrifugation, nuclei were harvested from the 25/75 interface. This nuclear material was re-loaded over 25% Percoll and centrifuged again at 2500 × g for 5 minutes. The supernatant was discarded, and the pellet was washed with 1mL Buffer B, typically yielding 100k nuclei per 0.1g tissue. For ATAC-Seq library preparation, each biological sample utilized 50k nuclei. Libraries were prepared using the Hyperactive ATAC-Seq Library Prep Kit (Vazyme TD711-01) and sequenced by Azenta or Novogene.

### ATAC-Seq analysis

For the ATAC-Seq analysis, the process commenced with the quality trimming of raw .fastq files using fastp, followed by alignment to the *Arabidopsis thaliana* TAIR10 reference genome via Bowtie2. The resultant BAM files were then filtered with Sambamba view to exclude unmapped reads, secondary alignments, and reads not meeting quality control criteria. Duplicates were marked using Sambamba markdup, and SAMtool view was employed to further eliminate low-quality reads. These processed BAM files were intersected with reference BED files using BEDTools. the BAM files were converted to bigwig using bamCoverage provided by deepTools v3.1.2 with a bin size of 10 bp and normalized by Bin Per Million mapped reads. Peak calling was performed using MACS2, with normalization and quantification undertaken by DiffBind. ChIPseeker was used for the annotation of these peaks. For Gene Ontology (GO) analysis of the enriched peaks, clusterProfiler was applied. Additionally, deepTools was utilized to assess TSS enrichment.

Annotated peaks were integrated with Weighted Gene Co-expression Network Analysis (WGCNA) results to perform Transcription Factor Binding Site (TFBS) enrichment analysis. The TFBS data were sourced from PlantTFDB v5.0 (Tian et al., 2020). For each WGCNA module/eigengene, TFBS enrichment was conducted using the MEME suite’s Analysis of Motif Enrichment (AME). diffTF was employed to categorize the transcription factors (TFs) as activators or repressors, based on a comparative analysis of RNA-Seq and ATAC-Seq data for *DM1-DM2d* and *DM6-DM7*, against their respective WT controls. Additionally, ChromVAR was utilized to examine synergistic interactions and correlations among TFBS (Schep et al., 2017). The network of TFs and WGCNA modules was visualized using Cytoscape.

TOBIAS footprinting was performed to compare the putative binding events between *DMs* and Col-0 (Bentsen et al., 2020).

## Contributions

**Conceptualisation:** E.C. and D.H.; **Methodology:** D.H., J.W., and E.C.; **Investigation:** D.H. (line generation, transcriptomic analysis, ATAC-seq analysis, data integration, bioinformatics), J.W. (line generation, *adr1s* knockout construction), Z.S. (sequencing support) **Resources:** W.Z. (provided wz189 *DM6-DM7* line), E.C. (supervision, funding); **Writing—Original Draft:** D.H.; **Writing—Review & Editing:** D.H., R.R.Q.L., and E.C.; **Visualisation:** D.H., J.W. and R.R.Q.L.; **Supervision:** E.C.; **Project Administration:** E.C.; **Funding Acquisition:** E.C.

## Acknowledgments

The study was supported by Singapore National Research Foundation under its Competitive Research Programme (NRF-CRP22-2019-0001) and Singapore Food Agency (SFS_RND_SUFP_002_04).

## References

Albert, I., Böhm, H., Albert, M., Feiler, C. E., Imkampe, J., Wallmeroth, N., Brancato, C., Raaymakers, T. M., Oome, S., Zhang, H., et al. (2015). An RLP23-SOBIR1-BAK1 complex mediates NLP-triggered immunity. Nat Plants 1:15140.

Alcázar, R., García, A. V., Parker, J. E., and Reymond, M. (2009). Incremental steps toward incompatibility revealed by Arabidopsis epistatic interactions modulating salicylic acid pathway activation. Proceedings of the National Academy of Sciences 106:334–339.

Andolfo, G., Villano, C., Errico, A., Frusciante, L., Carputo, D., Aversano, R., and Ercolano, M. R. (2019). Inferring RPW8-NLRs’s evolution patterns in seed plants: case study in Vitis vinifera. Planta 251:32.

Asai, T., Tena, G., Plotnikova, J., Willmann, M. R., Chiu, W.-L., Gomez-Gomez, L., Boller, T., Ausubel, F. M., and Sheen, J. (2002). MAP kinase signalling cascade in Arabidopsis innate immunity. Nature 415:977–983.

Atanasov, K. E., Liu, C., Erban, A., Kopka, J., Parker, J. E., and Alcázar, R. (2018). NLR mutations suppressing immune hybrid incompatibility and their effects on disease resistance. Plant Physiology 177:1152–1169.

Aufsatz, W., Amry, D., and Grimm, C. (1998). The ECS1 gene of Arabidopsis encodes a plant cell wall-associated protein and is potentially linked to a locus influencing resistance to Xanthomonas campestris. Plant Mol Biol 38:965–976.

Baile, F., Merini, W., Hidalgo, I., and Calonje, M. (2021). EAR domain-containing transcription factors trigger PRC2-mediated chromatin marking in Arabidopsis. The Plant Cell 33:2701–2715.

Barragan, C. A., Wu, R., Kim, S.-T., Xi, W., Habring, A., Hagmann, J., Van de Weyer, A.-L., Zaidem, M., Ho, W. W. H., Wang, G., et al. (2019). RPW8/HR repeats control NLR activation in Arabidopsis thaliana. PLOS Genetics 15:e1008313–e1008313.

Barragan, A. C., Collenberg, M., Wang, J., Lee, R. R. Q., Cher, W. Y., Rabanal, F. A., Ashkenazy, H., Weigel, D., and Chae, E. (2021). A Truncated Singleton NLR Causes Hybrid Necrosis in Arabidopsis thaliana. Molecular Biology and Evolution 38:557–574.

Bent, A. F., Kunkel, B. N., Dahlbeck, D., Brown, K. L., Schmidt, R., Giraudat, J., Leung, J., and Staskawicz, B. J. (1994). RPS2 of Arabidopsis thaliana: a Leucine-Rich Repeat Class of Plant Disease Resistance Genes. Science 265:1856–1860.

Bentsen, M., Goymann, P., Schultheis, H., Klee, K., Petrova, A., Wiegandt, R., Fust, A., Preussner, J., Kuenne, C., Braun, T., et al. (2020). ATAC-seq footprinting unravels kinetics of transcription factor binding during zygotic genome activation. Nat Commun 11:4267.

Berest, I., Arnold, C., Reyes-Palomares, A., Palla, G., Rasmussen, K. D., Giles, H., Bruch, P.-M., Huber, W., Dietrich, S., Helin, K., et al. (2019). Quantification of Differential Transcription Factor Activity and Multiomics-Based Classification into Activators and Repressors: diffTF. Cell Reports 29:3147–3159.e12.

Bi, G., Su, M., Li, N., Liang, Y., Dang, S., Xu, J., Hu, M., Wang, J., Zou, M., Deng, Y., et al. (2021). The ZAR1 resistosome is a calcium-permeable channel triggering plant immune signaling. Cell Advance Access published May 2021, doi:10.1016/j.cell.2021.05.003.

Bischoff, V., Selbig, J., and Scheible, W.-R. (2010). Involvement of TBL/DUF231 proteins into cell wall biology. Plant Signaling & Behavior 5:1057–1059.

Bomblies, K., and Weigel, D. (2007). Hybrid necrosis: autoimmunity as a potential gene-flow barrier in plant species. Nat Rev Genet 8:382–393.

Bomblies, K., Lempe, J., Epple, P., Warthmann, N., Lanz, C., Dangl, J. L., and Weigel, D. (2007). Autoimmune response as a mechanism for a Dobzhansky-Muller-type incompatibility syndrome in plants. PLoS Biology 5:1962–1972.

Bonardi, V., Tang, S., Stallmann, A., Roberts, M., Cherkis, K., and Dangl, J. L. (2011). Expanded functions for a family of plant intracellular immune receptors beyond specific recognition of pathogen effectors. Proceedings of the National Academy of Sciences 108:16463–16468.

Boutrot, F., and Zipfel, C. (2017). Function, Discovery, and Exploitation of Plant Pattern Recognition Receptors for Broad-Spectrum Disease Resistance. Annu. Rev. Phytopathol. 55:257–286.

Brooks, D. M., Bender, C. L., and Kunkel, B. N. (2005). The Pseudomonas syringae phytotoxin coronatine promotes virulence by overcoming salicylic acid-dependent defences in Arabidopsis thaliana. Molecular Plant Pathology 6:629–639.

Buscaill, P., and Rivas, S. (2014). Transcriptional control of plant defence responses. Current Opinion in Plant Biology 20:35–46.

Chae, E., Bomblies, K., Kim, S., Karelina, D., Zaidem, M., Ossowski, S., Martín-pizarro, C., Laitinen, R. A. E., Rowan, B. A., Tenenboim, H., et al. (2014). Species-wide Genetic Incompatibility Analysis Identifies Immune Genes as Hotspots of Deleterious Epistasis 159:1341–1351.

Chen, J., Zhang, X., Rathjen, J. P., and Dodds, P. N. (2022). Direct recognition of pathogen effectors by plant NLR immune receptors and downstream signalling. Essays in Biochemistry 66:471–483.

Chen, J., Li, L., Kim, J. H., Neuhäuser, B., Wang, M., Thelen, M., Hilleary, R., Chi, Y., Wei, L., Venkataramani, K., et al. (2023). Small proteins modulate ion-channel-like ACD6 to regulate immunity in Arabidopsis thaliana. Molecular Cell 83:4386–4397.e9.

Chhillar, H., Nguyen, H. H., Yeh, P.-M., Jones, J. D. G., and Ding, P. (2025). Modular mechanisms of immune priming and growth inhibition mediated by plant effector-triggered immunity. Cell Reports 44.

Cui, H., Tsuda, K., and Parker, J. E. (2015). Effector-triggered immunity: from pathogen perception to robust defense. Annu Rev Plant Biol 66:487–511.

Dangl, J. L., and Jones, J. D. G. (2001). Plant pathogens and integrated defence responses to infection. Nature 411:826–833.

de Oliveira, M. V. V., Xu, G., Li, B., de Souza Vespoli, L., Meng, X., Chen, X., Yu, X., de Souza, S. A., Intorne, A. C., de A. Manhães, A. M. E., et al. (2016). Specific control of Arabidopsis BAK1/SERK4-regulated cell death by protein glycosylation. Nat Plants 2:15218.

de Souza, A., Hull, P. A., Gille, S., and Pauly, M. (2014). Identification and functional characterization of the distinct plant pectin esterases PAE8 and PAE9 and their deletion mutants. Planta 240:1123–1138.

Ding, B., and Wang, G.-L. (2015). Chromatin versus pathogens: the function of epigenetics in plant immunity. Front. Plant Sci. 6.

Dong, O. X., Tong, M., Bonardi, V., Kasmi, F. E., Woloshen, V., Wünsch, L. K., Dangl, J. L., and Li, X. (2016). TNL-mediated immunity in Arabidopsis requires complex regulation of the redundant ADR1 gene family. New Phytologist 210:960–973.

Dongus, J. A., and Parker, J. E. (2021). EDS1 signalling: At the nexus of intracellular and surface receptor immunity. Current Opinion in Plant Biology 62:102039.

Feng, B., Han, Y., Xiao, Y., Kuang, J., Fan, Z., Chen, J., and Lu, W. (2016). The banana fruit Dof transcription factor MaDof23 acts as a repressor and interacts with MaERF9 in regulating ripening-related genes. J Exp Bot 67:2263–2275.

Förderer, A., Yu, D., Li, E., and Chai, J. (2022a). Resistosomes at the interface of pathogens and plants. Current Opinion in Plant Biology 67:102212.

Förderer, A., Li, E., Lawson, A. W., Deng, Y., Sun, Y., Logemann, E., Zhang, X., Wen, J., Han, Z., Chang, J., et al. (2022b). A wheat resistosome defines common principles of immune receptor channels. Nature Advance Access published September 26, 2022, doi:10.1038/s41586-022-05231-w.

Fornara, F., Panigrahi, K. C. S., Gissot, L., Sauerbrunn, N., Rühl, M., Jarillo, J. A., and Coupland, G. (2009). Arabidopsis DOF Transcription Factors Act Redundantly to Reduce CONSTANS Expression and Are Essential for a Photoperiodic Flowering Response. Developmental Cell 17:75–86.

Grant, J. J., Chini, A., Basu, D., and Loake, G. J. (2003). Targeted Activation Tagging of the Arabidopsis NBS-LRR gene, ADR1, Conveys Resistance to Virulent Pathogens. MPMI 16:669–680.

Guo, Y.-L., Fitz, J., Schneeberger, K., Ossowski, S., Cao, J., and Weigel, D. (2011). Genome-Wide Comparison of Nucleotide-Binding Site-Leucine-Rich Repeat-Encoding Genes in Arabidopsis. Plant Physiology 157:757–769.

He, Z., Webster, S., and He, S. Y. (2022). Growth–defense trade-offs in plants. Current Biology 32:R634–R639.

Henriques, R., Wang, H., Liu, J., Boix, M., Huang, L.-F., and Chua, N.-H. (2017). The antiphasic regulatory module comprising CDF5 and its antisense RNA FLORE links the circadian clock to photoperiodic flowering. New Phytologist 216:854–867.

Hinsch, M., and Staskawicz, B. (1996). Identification of a new Arabidopsis disease resistance locus, RPs4, and cloning of the corresponding avirulence gene, avrRps4, from Pseudomonas syringae pv. pisi. Mol Plant Microbe Interact 9:55–61.

Horsefield, S., Burdett, H., Zhang, X., Manik, M. K., Shi, Y., Chen, J., Qi, T., Gilley, J., Lai, J.-S., Rank, M. X., et al. (2019). NAD+ cleavage activity by animal and plant TIR domains in cell death pathways. Science 365:793–799.

Hu, Y., Dong, Q., and Yu, D. (2012). Arabidopsis WRKY46 coordinates with WRKY70 and WRKY53 in basal resistance against pathogen Pseudomonas syringae. Plant Sci 185–186:288–297.

Hu, C., Zhu, Y., Cui, Y., Cheng, K., Liang, W., Wei, Z., Zhu, M., Yin, H., Zeng, L., Xiao, Y., et al. (2018). A group of receptor kinases are essential for CLAVATA signalling to maintain stem cell homeostasis. Nature Plants 4:205–211.

Huang, S., Jia, A., Song, W., Hessler, G., Meng, Y., Sun, Y., Xu, L., Laessle, H., Jirschitzka, J., Ma, S., et al. (2022). Identification and receptor mechanism of TIR-catalyzed small molecules in plant immunity. Science 377:eabq3297.

Huot, B., Yao, J., Montgomery, B. L., and He, S. Y. (2014). Growth–Defense Tradeoffs in Plants: A Balancing Act to Optimize Fitness. Molecular Plant 7:1267–1287.

Ishiga, Y., Ishiga, T., Uppalapati, S. R., and Mysore, K. S. (2011). Arabidopsis seedling flood-inoculation technique: a rapid and reliable assay for studying plant-bacterial interactions. Plant Methods 7:32.

Ishiga, Y., Ishiga, T., Ichinose, Y., and Mysore, K. (2017). Pseudomonas syringae Flood-inoculation Method in Arabidopsis. BIO-PROTOCOL 7.

Jacob, F., Kracher, B., Mine, A., Seyfferth, C., Blanvillain-Baufumé, S., Parker, J. E., Tsuda, K., Schulze-Lefert, P., and Maekawa, T. (2018). A dominant-interfering camta3 mutation compromises primary transcriptional outputs mediated by both cell surface and intracellular immune receptors in Arabidopsis thaliana. New Phytologist 217:1667–1680.

Jacob, P., Hige, J., and Dangl, J. L. (2023). Is localized acquired resistance the mechanism for effector-triggered disease resistance in plants? Nat. Plants Advance Access published August 3, 2023, doi:10.1038/s41477-023-01466-1.

Jia, M., Shen, X., Tang, Y., Shi, X., and Gu, Y. (2021). A karyopherin constrains nuclear activity of the NLR protein SNC1 and is essential to prevent autoimmunity in Arabidopsis. Molecular Plant 14:1733–1744.

Jia, A., Huang, S., Song, W., Wang, J., Meng, Y., Sun, Y., Xu, L., Laessle, H., Jirschitzka, J., Hou, J., et al. (2022). TIR-catalyzed ADP-ribosylation reactions produce signaling molecules for plant immunity. Science 377:eabq8180.

Kim, N. H., Jacob, P., and Dangl, J. L. (2022). Con-Ca2+ -tenating plant immune responses via calcium-permeable cation channels. New Phytol 234:813–818.

Kumar, D., Kumar, R., Baek, D., Hyun, T.-K., Chung, W. S., Yun, D.-J., and Kim, J.-Y. (2017). Arabidopsis thaliana RECEPTOR DEAD KINASE1 Functions as a Positive Regulator in Plant Responses to ABA. Molecular Plant 10:223–243.

Langfelder, P., and Horvath, S. (2008). WGCNA: an R package for weighted correlation network analysis. BMC Bioinformatics 9:559.

Lapin, D., Kovacova, V., Sun, X., Dongus, J. A., Bhandari, D., von Born, P., Bautor, J., Guarneri, N., Rzemieniewski, J., Stuttmann, J., et al. (2019). A Coevolved EDS1-SAG101-NRG1 Module Mediates Cell Death Signaling by TIR-Domain Immune Receptors. The Plant Cell 31:2430–2455.

Lapin, D., Bhandari, D. D., and Parker, J. E. (2020). Origins and Immunity Networking Functions of EDS1 Family Proteins. Annual Review of Phytopathology 58:253–276.

Lee, R. R. Q., and Chae, E. (2025). Monkeys at Rigged Typewriters: A Population and Network View of Plant Immune System Incompatibility. Annual Review of Plant Biology 76:523–550.

Li, L., Habring, A., Wang, K., and Weigel, D. (2020). Atypical Resistance Protein RPW8/HR Triggers Oligomerization of the NLR Immune Receptor RPP7 and Autoimmunity. Cell Host and Microbe 27:405–417.e6.

Liu, Y., Zeng, Z., Zhang, Y.-M., Li, Q., Jiang, X.-M., Jiang, Z., Tang, J.-H., Chen, D., Wang, Q., Chen, J.-Q., et al. (2021). An angiosperm NLR Atlas reveals that NLR gene reduction is associated with ecological specialization and signal transduction component deletion. Molecular Plant 14:2015–2031.

Lu, H., Rate, D. N., Song, J. T., and Greenberg, J. T. (2003a). ACD6, a Novel Ankyrin Protein, Is a Regulator and an Effector of Salicylic Acid Signaling in the Arabidopsis Defense Response. Plant Cell 15:2408–2420.

Lu, H., Rate, D. N., Song, J. T., and Greenberg, J. T. (2003b). ACD6, a Novel Ankyrin Protein, Is a Regulator and an Effector of Salicylic Acid Signaling in the Arabidopsis Defense Response. Plant Cell 15:2408–2420.

Luo, M., Liu, X., Singh, P., Cui, Y., Zimmerli, L., and Wu, K. (2012). Chromatin modifications and remodeling in plant abiotic stress responses. Biochimica et Biophysica Acta (BBA) - Gene Regulatory Mechanisms 1819:129–136.

Ngou, B. P. M., Ahn, H.-K., Ding, P., and Jones, J. D. G. (2021). Mutual potentiation of plant immunity by cell-surface and intracellular receptors. Nature Advance Access published March 10, 2021, doi:10.1038/s41586-021-03315-7.

Nobori, T., Monell, A., Lee, T. A., Sakata, Y., Shirahama, S., Zhou, J., Nery, J. R., Mine, A., and Ecker, J. R. (2025). A rare PRIMER cell state in plant immunity. Nature Advance Access published January 8, 2025, doi:10.1038/s41586-024-08383-z.

Park, D. H., Lim, P. O., Kim, J. S., Cho, D. S., Hong, S. H., and Nam, H. G. (2003). The Arabidopsis COG1 gene encodes a Dof domain transcription factor and negatively regulates phytochrome signaling. The Plant Journal 34:161–171.

Pruitt, R. N., Locci, F., Wanke, F., Zhang, L., Saile, S. C., Joe, A., Karelina, D., Hua, C., Fröhlich, K., Wan, W.-L., et al. (2021). The EDS1–PAD4–ADR1 node mediates Arabidopsis pattern-triggered immunity. Nature Advance Access published September 8, 2021, doi:10.1038/s41586-021-03829-0.

Qin, H., Cheng, J., Han, G., and Gong, Z. (2024). Phylogenomic insights into the diversity and evolution of RPW8 - NLRS and their partners in plants. The Plant Journal 120:1032–1046.

Ramirez-Prado, J. S., Abulfaraj, A. A., Rayapuram, N., Benhamed, M., and Hirt, H. (2018). Plant Immunity: From Signaling to Epigenetic Control of Defense. Trends in Plant Science 23:833–844.

Rich-Griffin, C., Eichmann, R., Reitz, M. U., Hermann, S., Woolley-Allen, K., Brown, P. E., Wiwatdirekkul, K., Esteban, E., Pasha, A., Kogel, K.-H., et al. (2020). Regulation of Cell Type-Specific Immunity Networks in Arabidopsis Roots[CC-BY]. Plant Cell 32:2742–2762.

Rodriguez-Furlan, C., Campos, R., Toth, J. N., and Van Norman, J. M. (2022). Distinct mechanisms orchestrate the contra-polarity of IRK and KOIN, two LRR-receptor-kinases controlling root cell division. Nat Commun 13:235.

Saijo, Y., and Loo, E. P. (2020). Plant immunity in signal integration between biotic and abiotic stress responses. New Phytologist 225:87–104.

Saijo, Y., Loo, E. P., and Yasuda, S. (2018). Pattern recognition receptors and signaling in plant–microbe interactions. The Plant Journal 93:592–613.

Saile, S. C., Jacob, P., Castel, B., Jubic, L. M., Salas-Gonzáles, I., Bäcker, M., Jones, J. D. G., Dangl, J. L., and El Kasmi, F. (2020). Two unequally redundant “helper” immune receptor families mediate Arabidopsis thaliana intracellular “sensor” immune receptor functions. PLoS Biol 18:e3000783.

Schep, A. N., Wu, B., Buenrostro, J. D., and Greenleaf, W. J. (2017). chromVAR: inferring transcription-factor-associated accessibility from single-cell epigenomic data. Nat Methods 14:975–978.

Schulze, S., Yu, L., Hua, C., Zhang, L., Kolb, D., Weber, H., Ehinger, A., Saile, S. C., Stahl, M., Franz-Wachtel, M., et al. (2022). The *Arabidopsis* TIR-NBS-LRR protein CSA1 guards BAK1-BIR3 homeostasis and mediates convergence of pattern- and effector-induced immune responses. Cell Host & Microbe 30:1717–1731.e6.

Shpak, E. D., Berthiaume, C. T., Hill, E. J., and Torii, K. U. (2004). Synergistic interaction of three ERECTA-family receptor-like kinases controls Arabidopsis organ growth and flower development by promoting cell proliferation. Development 131:1491–1501.

Steidele, C. E., and Stam, R. (2021). Multi-omics approach highlights differences between RLP classes in Arabidopsis thaliana. BMC Genomics 22:1–14.

Świadek, M., Proost, S., Sieh, D., Yu, J., Todesco, M., Jorzig, C., Rodriguez Cubillos, A. E., Plötner, B., Nikoloski, Z., Chae, E., et al. (2017). Novel allelic variants in ACD6 cause hybrid necrosis in local collection of Arabidopsis thaliana. New Phytologist 213:900–915.

Thomma, B. P. H. J., Nürnberger, T., and Joosten, M. H. A. J. (2011). Of PAMPs and effectors: the blurred PTI-ETI dichotomy. Plant Cell 23:4–15.

Tian, H., Wu, Z., Chen, S., Ao, K., Huang, W., Yaghmaiean, H., Sun, T., Xu, F., Zhang, Y., Wang, S., et al. (2021). Activation of TIR signalling boosts pattern-triggered immunity. Nature 598:500–503.

Todesco, M., Balasubramanian, S., Hu, T. T., Traw, M. B., Horton, M., Epple, P., Kuhns, C., Sureshkumar, S., Schwartz, C., Lanz, C., et al. (2010). Natural allelic variation underlying a major fitness trade-off in Arabidopsis thaliana. Nature 465:632–636.

Todesco, M., Kim, S.-T., Chae, E., Bomblies, K., Zaidem, M., Smith, L. M., Weigel, D., and Laitinen, R. A. E. (2014). Activation of the Arabidopsis thaliana Immune System by Combinations of Common ACD6 Alleles. PLOS Genetics 10:e1004459.

Tran, D. T. N., Chung, E. H., Habring-Müller, A., Demar, M., Schwab, R., Dangl, J. L., Weigel, D., and Chae, E. (2017). Activation of a Plant NLR Complex through Heteromeric Association with an Autoimmune Risk Variant of Another NLR. Current Biology 27:1148–1160.

Tsuda, K., and Katagiri, F. (2010). Comparing signaling mechanisms engaged in pattern-triggered and effector-triggered immunity. Current Opinion in Plant Biology 13:459–465.

Tsuda, K., and Somssich, I. E. (2015). Transcriptional networks in plant immunity. New Phytologist 206:932–947.

Tsuda, K., Sato, M., Stoddard, T., Glazebrook, J., and Katagiri, F. (2009). Network Properties of Robust Immunity in Plants. PLOS Genetics 5:e1000772.

Urquhart, W., Chin, K., Ung, H., Moeder, W., and Yoshioka, K. (2011). The cyclic nucleotide-gated channels AtCNGC11 and 12 are involved in multiple Ca2+-dependent physiological responses and act in a synergistic manner. Journal of Experimental Botany 62:3671–3682.

Van de Weyer, A. L., Monteiro, F., Furzer, O. J., Nishimura, M. T., Cevik, V., Witek, K., Jones, J. D. G., Dangl, J. L., Weigel, D., and Bemm, F. (2019). A Species-Wide Inventory of NLR Genes and Alleles in Arabidopsis thaliana. Cell 178:1260–1272.e14.

van der Hoorn, R. A. L., and Kamoun, S. (2008). From Guard to Decoy: A New Model for Perception of Plant Pathogen Effectors. The Plant Cell 20:2009–2017.

Wan, L., Essuman, K., Anderson, R. G., Sasaki, Y., Monteiro, F., Chung, E.-H., Osborne Nishimura, E., DiAntonio, A., Milbrandt, J., Dangl, J. L., et al. (2019). TIR domains of plant immune receptors are NAD ^+^ -cleaving enzymes that promote cell death. Science 365:799–803.

Wang, W., Liu, N., Gao, C., Rui, L., Jiang, Q., Chen, S., Zhang, Q., Zhong, G., and Tang, D. (2021). The truncated TNL receptor TN2-mediated immune responses require ADR1 function. Plant J 108:672–689.

Wei, P.-C., Tan, F., Gao, X.-Q., Zhang, X.-Q., Wang, G.-Q., Xu, H., Li, L.-J., Chen, J., and Wang, X.-C. (2010). Overexpression of AtDOF4.7, an Arabidopsis DOF Family Transcription Factor, Induces Floral Organ Abscission Deficiency in Arabidopsis1[C][W]. Plant Physiol 153:1031–1045.

Wu, Z., Tian, L., Liu, X., Zhang, Y., and Li, X. (2021). TIR signal promotes interactions between lipase-like proteins and ADR1-L1 receptor and ADR1-L1 oligomerization. Plant Physiol 187:681–686.

Yang, W., Dong, R., Liu, L., Hu, Z., Li, J., Wang, Y., Ding, X., and Chu, Z. (2016). A novel mutant allele of SSI2 confers a better balance between disease resistance and plant growth inhibition on Arabidopsis thaliana. BMC Plant Biology 16:208.

Yang, L., Chen, X., Wang, Z., Sun, Q., Hong, A., Zhang, A., Zhong, X., and Hua, J. (2020). HOS15 and HDA9 negatively regulate immunity through histone deacetylation of intracellular immune receptor NLR genes in Arabidopsis. New Phytologist 226:507–522.

Yang, L., Wang, Z., and Hua, J. (2021). A Meta-Analysis Reveals Opposite Effects of Biotic and Abiotic Stresses on Transcript Levels of Arabidopsis Intracellular Immune Receptor Genes. Front. Plant Sci. 12.

Yang, Y., Kim, N. H., Cevik, V., Jacob, P., Wan, L., Furzer, O. J., and Dangl, J. L. (2022). Allelic variation in the *Arabidopsis* TNL CHS3/CSA1 immune receptor pair reveals two functional cell-death regulatory modes. Cell Host & Microbe 30:1701–1716.e5.

Yuan, M., Jiang, Z., Bi, G., Nomura, K., Liu, M., Wang, Y., Cai, B., Zhou, J.-M., He, S. Y., and Xin, X.-F. (2021). Pattern-recognition receptors are required for NLR-mediated plant immunity. Nature Advance Access published March 10, 2021, doi:10.1038/s41586-021-03316-6.

Zeilmaker, T., Ludwig, N. R., Elberse, J., Seidl, M. F., Berke, L., Van Doorn, A., Schuurink, R. C., Snel, B., and Van den Ackerveken, G. (2015). DOWNY MILDEW RESISTANT 6 and DMR6-LIKE OXYGENASE 1 are partially redundant but distinct suppressors of immunity in Arabidopsis. The Plant Journal 81:210–222.

Zhang, Z., Liu, Y., Ding, P., Li, Y., Kong, Q., and Zhang, Y. (2014). Splicing of Receptor-Like Kinase-Encoding SNC4 and CERK1 is Regulated by Two Conserved Splicing Factors that Are Required for Plant Immunity. Molecular Plant 7:1766–1775.

Zhang, C., Gao, M., Seitz, N. C., Angel, W., Hallworth, A., Wiratan, L., Darwish, O., Alkharouf, N., Dawit, T., Lin, D., et al. (2019). LUX ARRHYTHMO mediates crosstalk between the circadian clock and defense in Arabidopsis. Nat Commun 10:2543.

Zheng, X., Spivey, N. W., Zeng, W., Liu, P.-P., Fu, Z. Q., Klessig, D. F., He, S. Y., and Dong, X. (2012). Coronatine promotes Pseudomonas syringae virulence in plants by activating a signaling cascade that inhibits salicylic acid accumulation. Cell host & microbe 11:587.

Zhu, W., Zaidem, M., Weyer, A.-L. V. de, Gutaker, R. M., Chae, E., Kim, S.-T., Bemm, F., Li, L., Todesco, M., Schwab, R., et al. (2018). Modulation of ACD6 dependent hyperimmunity by natural alleles of an Arabidopsis thaliana NLR resistance gene. PLOS Genetics 14:e1007628.

Zou, X., and Sun, H. (2023). DOF transcription factors: Specific regulators of plant biological processes. Frontiers in Plant Science 14.

